# Evolution of enhanced innate immune evasion by the SARS-CoV-2 B.1.1.7 UK variant

**DOI:** 10.1101/2021.06.06.446826

**Authors:** Lucy G Thorne, Mehdi Bouhaddou, Ann-Kathrin Reuschl, Lorena Zuliani-Alvarez, Ben Polacco, Adrian Pelin, Jyoti Batra, Matthew V.X. Whelan, Manisha Ummadi, Ajda Rojc, Jane Turner, Kirsten Obernier, Hannes Braberg, Margaret Soucheray, Alicia Richards, Kuei-Ho Chen, Bhavya Harjai, Danish Memon, Myra Hosmillo, Joseph Hiatt, Aminu Jahun, Ian G. Goodfellow, Jacqueline M. Fabius, Kevan Shokat, Natalia Jura, Klim Verba, Mahdad Noursadeghi, Pedro Beltrao, Danielle L. Swaney, Adolfo Garcia-Sastre, Clare Jolly, Greg J. Towers, Nevan J. Krogan

## Abstract

Emergence of SARS-CoV-2 variants, including the globally successful B.1.1.7 lineage, suggests viral adaptations to host selective pressures resulting in more efficient transmission. Although much effort has focused on Spike adaptation for viral entry and adaptive immune escape, B.1.1.7 mutations outside Spike likely contribute to enhance transmission. Here we used unbiased abundance proteomics, phosphoproteomics, mRNA sequencing and viral replication assays to show that B.1.1.7 isolates more effectively suppress host innate immune responses in airway epithelial cells. We found that B.1.1.7 isolates have dramatically increased subgenomic RNA and protein levels of Orf9b and Orf6, both known innate immune antagonists. Expression of Orf9b alone suppressed the innate immune response through interaction with TOM70, a mitochondrial protein required for RNA sensing adaptor MAVS activation, and Orf9b binding and activity was regulated via phosphorylation. We conclude that B.1.1.7 has evolved beyond the Spike coding region to more effectively antagonise host innate immune responses through upregulation of specific subgenomic RNA synthesis and increased protein expression of key innate immune antagonists. We propose that more effective innate immune antagonism increases the likelihood of successful B.1.1.7 transmission, and may increase *in vivo* replication and duration of infection.

## Main

The SARS-CoV-2 B.1.1.7 lineage was detected in the United Kingdom in September 2020 and quickly became the dominant variant worldwide^1^. Epidemiologically, B.1.1.7 human-to-human transmission is superior to other SARS-CoV-2 lineages^2,3^, making it a variant of concern (VOC), threatening public health containment measures^4^. B.1.1.7 infection has been associated with enhanced clinical severity in the community in the UK, although a clear association with increased mortality has not yet emerged ^2,3,5,6^.

B.1.1.7 is defined by a constellation of 23 mutations^7^: 17 that alter protein sequence (14 non-synonymous mutations and 3 deletions) and 6 synonymous mutations (Fig. 1a). Protein coding changes concentrate in Spike, which facilitates viral entry through interaction with the human receptor ACE2^8^. This has led the field to focus on understanding viral escape from wave one (early-lineage) driven adaptive immunity and its implications for infection control and vaccine development. Fortunately, despite adaptation of Spike, B.1.1.7 remains sensitive to vaccine- and infection-induced neutralising antibodies^9–11^. B.1.1.7 variant of concern (VOC)-defining mutations outside Spike suggest that Spike-independent adaptation to host may contribute to the B.1.1.7 transmission advantage. Most B.1.1.7 coding changes map to non-structural proteins Nsp3, Nsp6, accessory protein Orf8 and nucleocapsid protein (N), all of which have been shown to modulate the innate immune response^12–16^. Furthermore, it is unclear whether any of the B.1.1.7-specific mutations impact the expression levels of viral proteins. In sum, the impact of these additional mutations on viral replication, transmission and pathogenesis has not been characterised.

**Figure 1.**
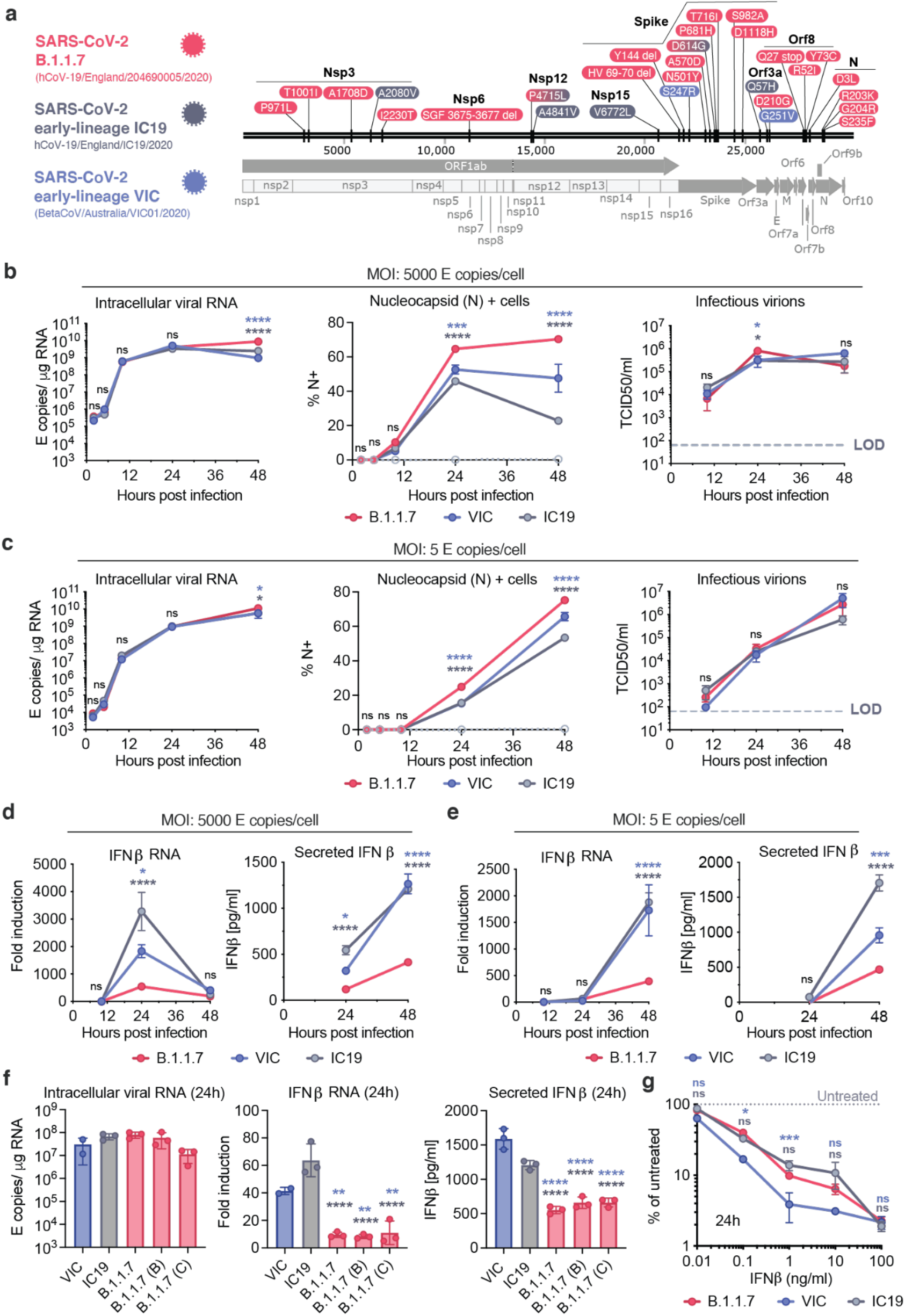
SARS-CoV-2 B.1.1.7 antagonises innate immune activation more efficiently than early-lineage isolates. **a.** SARS-CoV-2 viruses compared in this study. Protein coding changes in B.1.1.7 (red), IC19 (grey) and VIC (blue) are indicated in comparison to the Wuhan-Hu-1 reference genome (MN908947). B.1.1.7 changes include 23 lineage defining mutations, plus additional changes compared to Wuhan-Hu-1, totalling 29. **b** and **c.** Calu-3 cells were infected with either (b) 5000 E copies/cell or (c) 5 E copies/cell of B.1.1.7, VIC and IC19. Measurements of replication of SARS-CoV-2 genomic and subgenomic E RNAs (RT-qPCR) (left), % infection by intracellular nucleocapsid positivity (centre) or infectious virion production by TCID50/ml (right) over time are shown. **d** and **e.** Fold induction of IFNβ gene expression and protein secretion over time from cells in (b) and (c) respectively. **f.** Replication (24hpi), IFNβ induction (24hpi) and IFNβ secretion (48hpi) by multiple independent B.1.1.7 isolates compared to IC19 and VIC at 250 E copies/cell. **g.** SARS-CoV-2 infection at 2000 E copies/cell after 8h pre-treatment with IFNβ at the indicated concentrations. Infection is shown as intracellular N levels normalised to untreated controls at 24hpi. Data shown are mean +/- SEM of one of three representative experiments performed in triplicate. Statistical comparisons are performed by Two Way ANOVA (a,b,c,d,g) or One Way ANOVA with a Tukey post-comparison test (f). Blue stars indicate comparison between B.1.1.7 and VIC (blue lines and symbols), grey stars indicate comparison between B.1.1.7 and IC19 (grey lines and symbols). * (p<0.05), ** (p<0.01), *** (p<0.001), **** (p<0.0001). ns: non-significant. E: viral envelope gene. Hpi: hours post infection.

Innate immune responses can exert strong selective pressure during viral transmission ^17–19^ and play an important role in determining clinical outcomes to SARS-CoV-2 infection^20–22^. We therefore reasoned that B.1.1.7 may have evolved to enhance innate immune escape. We and others have recently shown that infection of naturally permissive Calu-3 human lung epithelial cells with a wave one SARS-CoV-2 lineage B isolate (BetaCoV/Australia/VIC01/2020, VIC) induces a robust but delayed innate response, driven by activation of RNA sensors RIG-I and MDA5 ^23^. A delayed response, compared to rapid viral RNA replication, suggests effective innate immune antagonism and evasion by SARS-CoV-2 early in infection^13,24^. Furthermore, the gene expression changes observed during late innate responses in infected Calu-3 cells reflect the overarching inflammatory signatures observed at the site of infection and those associated with severe COVID-19 ^25–28^. Here, we used the Calu-3 cell model to evaluate differences between B.1.1.7 and wave one SARS-CoV-2 viruses.

### Comparative analysis of virus replication kinetics and interferon induction

We compared replication and innate immune activation for B.1.1.7 and two first wave (early lineage) isolates, B lineage isolate BetaCoV/Australia/VIC01/2020 (VIC) and B.1.13 lineage isolate hCoV-19/England/IC19/2020 (IC19) (Fig. 1a) in Calu-3 lung epithelial cells. Input dose was normalised using viral genome copies measured by RT-qPCR for the envelope (E) coding region. We found that B.1.1.7 replication was comparable to both wave one isolates at high and low multiplicity of infection (MOI), measuring intracellular E copies, positivity for nucleocapsid protein and infectious virion production by TCID50 on Hela-ACE2 cells (Fig. 1b, 1c). We observed a small but significant increase in N positivity for B.1.1.7 (Fig. 1b, 1c), which we explain later in the context of differences in viral protein expression.

Identical replication of all three isolates enabled direct comparison of the innate immune response without differences in the amount of viral RNA produced, the principal pathogen associated molecular pattern (PAMP)^23^, being a confounding factor. We found that B.1.1.7 infection led to lower levels of IFNβ expression and secretion, at both high and low MOI (Fig. 1d, 1e). Similar replication, but reduced IFNβ induction by B.1.1.7 was confirmed with two additional independent B.1.1.7 isolates (Fig. 1f), suggesting consistent enhancement of innate immune antagonism, or evasion, for B.1.1.7 lineage isolates.

As IFN resistance correlates with enhanced transmission of other pandemic viruses^17,18^, we compared sensitivity to IFNβ inhibition of B.1.1.7 and first wave isolates. B.1.1.7 was consistently less sensitive to IFNβ pre-treatment over a wide dose range, compared to first wave isolate VIC (lineage B) (Fig. 1g), suggesting that B.1.1.7 infection not only induces less IFNβ (Fig. 1d, 1e) but that it is also less sensitive to its effects. Interestingly, wave one IC19 (B.1.13) showed a similar reduction in IFNβ sensitivity as B.1.1.7. This may be due to the shared Spike mutation D614G in IC19 and B.1.1.7, but not VIC, which is associated with enhanced transmissibility and increased entry efficiency^29–31^. Indeed, D614G has been associated with resistance to a range of Type I and III IFNs across several SARS-CoV-2 lineages, and contributes to the enhanced IFN-evasion of B.1.1.7^32^. Type I IFN restriction of SARS-CoV-2 is mediated in part by interferon induced membrane protein 2 (IFITM2) suppression of viral entry, and IFITM2 sensitivity is influenced by the Spike sequence^33,34^. We therefore focused on characterising the mechanism of enhanced antagonism of the innate response which was unique to the B.1.1.7 lineage.

### Global proteomic and genomic analyses reveal enhanced innate immune suppression by B.1.1.7

To compare cellular host responses to SARS-CoV-2 variants, we performed global mass spectrometry-based protein abundance and phosphorylation profiling (i.e. phosphoproteomics) as well as total RNAseq on infected Calu-3 cells at 10 and 24 hours post infection (hpi) (Fig. 2a, Table S1). The proteomic analysis was performed using a data-independent acquisition (DIA) approach, which decreases sample-to-sample and time point variability in peptide detection over the traditional data-dependent acquisition (DDA) mode, strengthening the comparative potential of these datasets (see Methods). Compared to mock infection, we observed robust changes in RNA abundance and protein phosphorylation after infection, with fewer changes at the level of protein abundance (Fig. S1a). After quality control data filtering was performed (see Methods), principal components analysis (PCA; Fig. S1b) and Pearson’s correlation (Fig. S1c) confirmed strong correlation between biological replicates, time points, and conditions. On average, we quantified 15,000-16,000 mRNA transcripts above background levels (Fig. S1d), 33,000-40,000 peptides mapping to 3,600-4,000 proteins for protein abundance (Fig. S1e), and 22,000-30,000 phosphorylated peptides mapping to 3,200-3,800 proteins for phosphoproteomics (Fig. S1f).

**Figure 2.**
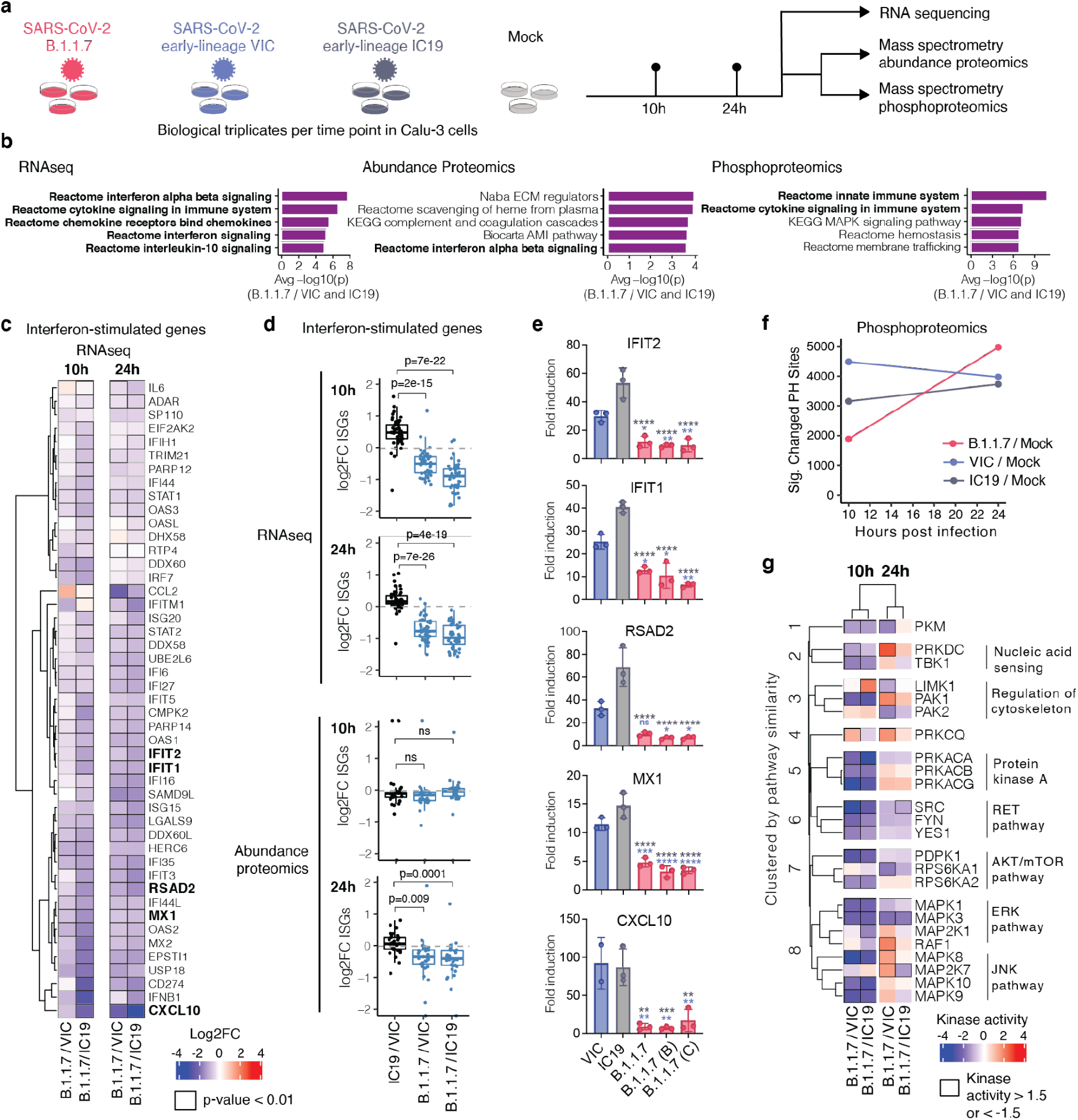
Global RNAseq and proteomics reveal innate immune suppression by B.1.1.7. **a.** Calu-3 cells were infected with SARS-CoV-2 B.1.1.7 (red) or early lineages VIC (blue) and IC19 (grey) at 5000 E copies/cell or mock-infected. At 10 and 24hpi, samples were harvested for phosphoproteomics and abundance proteomics analysis using a data-independent acquisition (DIA) approach. Separate wells were harvested for total RNA-sequencing. **b.** Unbiased pathway enrichment analysis was performed to compare B.1.1.7 to VIC and IC19 (see Methods). The −log10(p-values) were averaged for enrichments using B.1.1.7/VIC and B.1.1.7 /IC19 at 10 and 24hpi (4 data points total) and used to rank terms. The top 5 terms for each data type are displayed. Terms associated with the innate immune system are bolded. **c.** Heatmap depicting log2 fold change (color) of interferon-stimulated genes (ISGs)^36^ comparing B.1.1.7 to VIC or IC19 at 10 and 24hpi (see Methods[RAK1]). Squares outlined in black indicate a statistically significant fold change (p-value < 0.01). **d.** Box plots show log2 fold change of interferon stimulated genes (ISG) between B.1.1.7/VIC (blue), B.1.1.7/IC19 (blue) or IC19/VIC (black) in RNAseq and abundance proteomics dataset at 10 and 24hpi. Two-tailed student’s t-tests were performed for each comparison and p-values are displayed. **e.** Confirmatory RT-qPCR analysis of bolded ISGs from (a) expressed in Calu-3 cells infected with multiple B.1.1.7 isolates, VIC or IC19 at 2000 genomes/cell. **f.** The number of phosphorylation sites significantly dysregulated for B.1.1.7, VIC, or IC19 versus mock at 10 or 24hpi. Statistical significance was determined as absolute log2 FC > 1 and adjusted p-value < 0.05. **g.** Kinase activities for members of the top enriched terms for the phosphoproteomics dataset “Reactome innate immune system” (b, right), for each time point. Kinase activities were estimated from phosphoproteomics data using prior knowledge of kinase-substrate relationships. Kinases were clustered along the rows based on frequency of co-membership in pathway terms and manually annotated (see Methods). Data shown are mean +/- SEM (e). Statistical comparisons are performed by Two-tailed student’s t-tests (d) or Two Way ANOVA with a Tukey’s multiple comparisons post-test (e). Blue stars indicate comparison between B.1.1.7 and VIC (blue bars), grey stars indicate comparison between B.1.1.7 and IC19 (grey bars). * (p<0.05), ** (p<0.01), *** (p<0.001), **** (p<0.0001), or exact p-value are shown (d). ns: non-significant.

Gene set pathway enrichment^35^ analysis comparing B.1.1.7 to wave one isolates VIC and IC19 highlighted innate immune system-related pathways among the top 5 terms for all three data types (RNA, protein abundance, and phosphorylation) (Fig. 2b, S1g-i, Table S2). Top scoring terms were related to interferon alpha beta signalling and cytokine/chemokine signalling, and most predominantly enriched for the RNA and protein phosphorylation datasets (Fig. 2b). Concordantly, in addition to the reduction of IFNβ production (Fig 1d, 1e, 1f), B.1.1.7 infection resulted in reduced induction of interferon-stimulated genes (ISGs) measured in the RNAseq and protein abundance datasets using a predefined set of ISGs^36^ (detailed in the Methods, Table S3). This was evident at 10 and 24hpi at the RNA level (Fig. 2c–d, S2a, S2c) and at 24hpi for protein (Fig. 2d, S2b). For a subset of genes (*CXCL10*, *IFIT2*, *MX1*, *IFIT1*, and *RSAD2*), we confirmed reduced ISG induction by multiple B.1.1.7 isolates, compared to VIC and IC19 at 24hpi, using RT-qPCR (Fig. 2e).

Consistent with reduced innate immune activation by B.1.1.7, we observed lower overall changes in protein phosphorylation early in infection (10hpi) for B.1.1.7 compared to wave one isolates (Fig. 2f). Accordingly, gene set enrichment analysis revealed that the pathways highlighted by reduced phosphorylation at 10hpi are related to the innate immune response. These observations are indicative of enhanced innate immune antagonism by this variant. Strikingly, this was reversed at 24hpi as B.1.1.7 caused enhanced phosphorylation later in infection (Fig. S1i).

This notion of enhanced evasion at early time points, but increased activation at later time points by B.1.1.7 led us to investigate the differential regulation of kinase signalling cascades between and wave one viruses, especially in relation to innate immune signaling. We used the phosphoproteomics data to estimate kinase activities for 191 kinases based on regulation of their known substrates^37,38^ (Table S4), and grouped kinases according to their temporal dynamics (Fig. S2e). In a targeted approach, we compiled a list of kinases from the top enriched term (“Reactome innate immune system”; Fig. 2b) that were previously implicated in innate immune regulation and significantly dysregulated during infection. This identified 24 kinases, which we clustered by similar pathway membership (Fig. 2g and Methods). At 10hpi, we observed decreased activity of TBK1, a central kinase in nucleic acid sensing, as well as decreased activity in protein kinase A, PRKDC, RET, AKT/mTOR, ERK, and JNK pathways. Intriguingly, at 24hpi, TBK1, PRKDC, JNK, ERK, and PKA kinase activity was increased for B.1.1.7 compared to VIC (Fig. 2g), consistent with the increased phosphorylation in innate immune system enriched pathway terms (Fig. S1i). Thus, B.1.1.7 enhanced innate immune antagonism at the level of protein phosphorylation is only observed at early time points after infection suggesting a delay in the activation of the signalling pathways involved in viral recognition compared to early lineage viruses. However, later during infection as viral replication ramps up, it triggers the phosphorylation cascades leading to the activation of these pathways (Fig. 2f and Fig. S1i).

### B.1.1.7 has enhanced expression of subgenomic RNA and protein for key innate immune antagonists

We next examined the RNAseq and protein abundance mass spectrometry data of the viral genes and proteins seeking to further understand the differences between B.1.1.7 and wave one isolates that underlie the contrasting host responses (Fig. 3a, S3a, Table S6, Table S7). As RNA replication, measured by genomic and subgenomic E levels, was similar between variants (Fig. 1b, 1c), we determined the levels of each subgenomic RNA by selecting transcripts with a 5’ leader sequence, i.e. the segment derived from the 5’ genomic RNA during sgRNA transcription (Fig. 3a, S4). Importantly, we observed similar levels of viral Nsp1/2/3 protein translated from genomic RNA (Fig. 3a), which was again consistent with comparable levels of infection, enabling effective comparisons of transcription and protein expression between variants.

**Figure 3.**
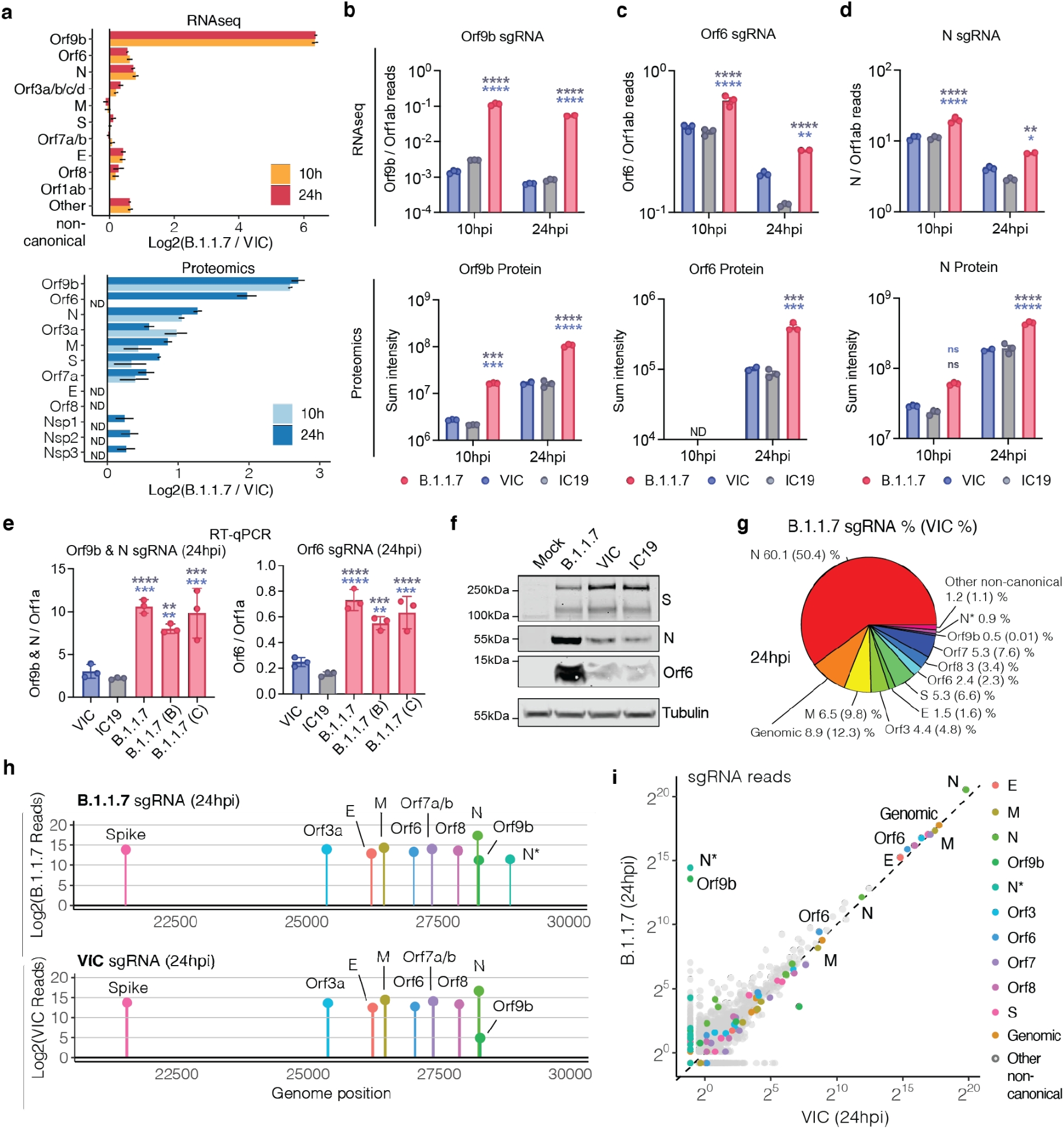
SARS-CoV-2 B.1.1.7 variant upregulates innate immune antagonists at the subgenomic RNA and protein level. **a.** Log2 ratio of B.1.1.7 to VIC subgenomic RNA (sgRNA) containing a leader sequence normalised to total genomic RNA per time point and virus (top). Log2 ratio of B.1.1.7 to VIC viral proteins quantified as determined from the abundance proteomics dataset (bottom). Peptide intensities are summed per viral protein. Only peptides detected in both B.1.1.7 and VIC are used for quantification. Bars depict the mean of three biological replicates. ND: not detected. **b. c.** and **d.** Quantification of Orf9b (b), Orf6 (c) and N (d) sgRNA from RNAseq dataset. Counts are normalised to genomic RNA abundance at each time point and virus (top). Bottom panels show summed peptides per viral protein from proteomics dataset (no normalisation). **e.** Quantification of Orf9b & N (left) or Orf6 (right) sgRNA abundance via RT-qPCR in independent B.1.1.7 isolates, VIC, or IC19. **f.** Western blot of Orf6, N and S expression in Calu-3 cells infected with B.1.1.7, VIC, or IC19 at 24hpi. **g.** Pie chart depicting proportion of total sgRNA mapping to each viral sgRNA (containing leader sequence) for B.1.1.7. VIC percentages in parentheses. **h.** sgRNA log2 normalised counts (dot height) at 24hpi for B.1.1.7 (top) or VIC (bottom) projected onto their identified start sites on the SARS-CoV-2 genome. Only canonical and two non-canonical sgRNAs (Orf9b and N*) are depicted. All other non-canonical sgRNAs were excluded. **i.** Scatter plot of sgRNA abundance in B.1.1.7 or VIC at 24hpi. Grey dots indicated other non-canonical sgRNAs containing a leader sequence but no clear start codon. For (a-e), mean +/- SEM are shown. Statistical comparisons for (c-e) were performed by Two Way ANOVA with Tukey’s multiple comparisons post-test. * (p<0.05), ** (p<0.01), *** (p<0.001), **** (p<0.0001). ns: non-significant.

Strikingly, we found a large (over 80-fold) increase in innate immune antagonist Orf9b sgRNA levels^39^, leading to a 6.5-fold increase in Orf9b protein levels for B.1.1.7 compared to VIC (Fig. 3a, 3b). Similarly, a 6.7-fold increase in protein and 64.5-fold increase in RNA was observed at 24hpi for B.1.1.7 compared to IC19 (Fig. S3a). Differential ORF9b expression was evident by 10hpi at the RNA and protein levels (Fig. 3a, 3b). The increase in B.1.1.7 Orf9b transcription might be attributable to the D3L mutation in N, which introduces an enhanced transcriptional regulator sequence (TRS) upstream of Orf9b, expressed as an alternative reading frame within N^40^. Alternatively, the D3L mutation close to the ATG of Orf9b might enhance its translation from the N sgRNA. In addition, when comparing B.1.1.7 to VIC, we found a significant but modest 1.5-fold increase in sgRNA levels for a second innate immune regulator, Orf6^13,24^ (2.1-fold compared to IC19). This corresponded to a 3.9-fold increase in Orf6 protein levels (4.6-fold compared to IC19) at 24hpi (Fig. 3a, 3c, Table S6).

Additionally, we detected elevated sgRNA levels in B.1.1.7 of a third innate immune regulator, nucleocapsid (N)^16^. B.1.1.7 N RNA was increased 1.7-fold and 2.3-fold compared to VIC and IC19, respectively, corresponding to a 2.4-fold and 2.3-fold increase in N protein levels (Fig. 3a, 3d). This increase in N might also be a contributor to the enhanced expression of Orf9b, as much of Orf9b is thought to be expressed from the same subgenomic RNA as the N protein. These results are consistent with the increase in N+ cells measured during Calu-3 infection (Fig. 1b,1c). We also observed enhancement of Orf3a, M, and Orf7b proteins at 24hpi for B.1.1.7, with only very modest changes observed at the RNA level (Fig. S3c,d). We confirmed upregulation of Orf9b, Orf6, N and Orf3a sgRNA using RT-qPCR (Fig. 3e, S3b) and confirmed heightened expression of B.1.1.7 Orf6 and N proteins by western blot (Fig. 3f). Unfortunately, we do not have a suitable ORF9b antibody for western blot. These findings are in line with the reported enhanced expression of B.1.1.7 sgRNA encoding Orf9b, Orf6, and N in clinical samples^40^. The proportion of each sgRNA of the total sgRNA reads are summarised for B.1.1.7 and wave one VIC in Fig. 3g and S3e. Intriguingly, we observed an additional sgRNA, N* ^40^, with an in-frame start codon at M210 encoding the C-terminal portion of N (Fig. 3h, Table S7), amounting for 0.9% of the total sgRNA for B.1.1.7 (Fig. 3g). We did not detect N* sgRNA in VIC or IC19 above background levels suggesting that the B.1.1.7 N R203K and R204K mutations, just upstream of the new N* start codon, may create a novel transcriptional regulatory sequence (TRS) permitting N* transcription, as previously hypothesised^40^. Indeed, sgRNA abundance measurements were consistent with Orf9b and N* being the most differentially expressed sgRNA between B.1.1.7 and VIC at 24hpi (Fig. 3i).

### Orf9b antagonises innate immune activation by interacting with human TOM70

To further understand the transcription pathways differentially activated by B.1.1.7, we used the RNAseq dataset to estimate transcription factor activities by mapping target genes to corresponding transcriptional regulators (Fig. S2d, Table S5). We extracted significantly regulated transcription factors within the top 5 most enriched terms from the unbiased RNAseq pathway enrichment analysis (Fig 2a, left). This revealed that IRF and STAT transcription factor families are significantly less activated by B.1.1.7 compared to wave one viruses (Fig. 4a). Consistently, measuring IRF3 nuclear translocation by single-cell immunofluorescence demonstrated reduced IRF3 activation after B.1.1.7 infection compared to VIC (Fig. 4b). STAT1/STAT2/IRF9 lie downstream of the Type I IFN receptor, and potent inhibition by B.1.1.7 is consistent with increased Orf6 levels, known to inhibit STAT1, and IRF3, nuclear translocation^13,24^.

**Figure 4.**
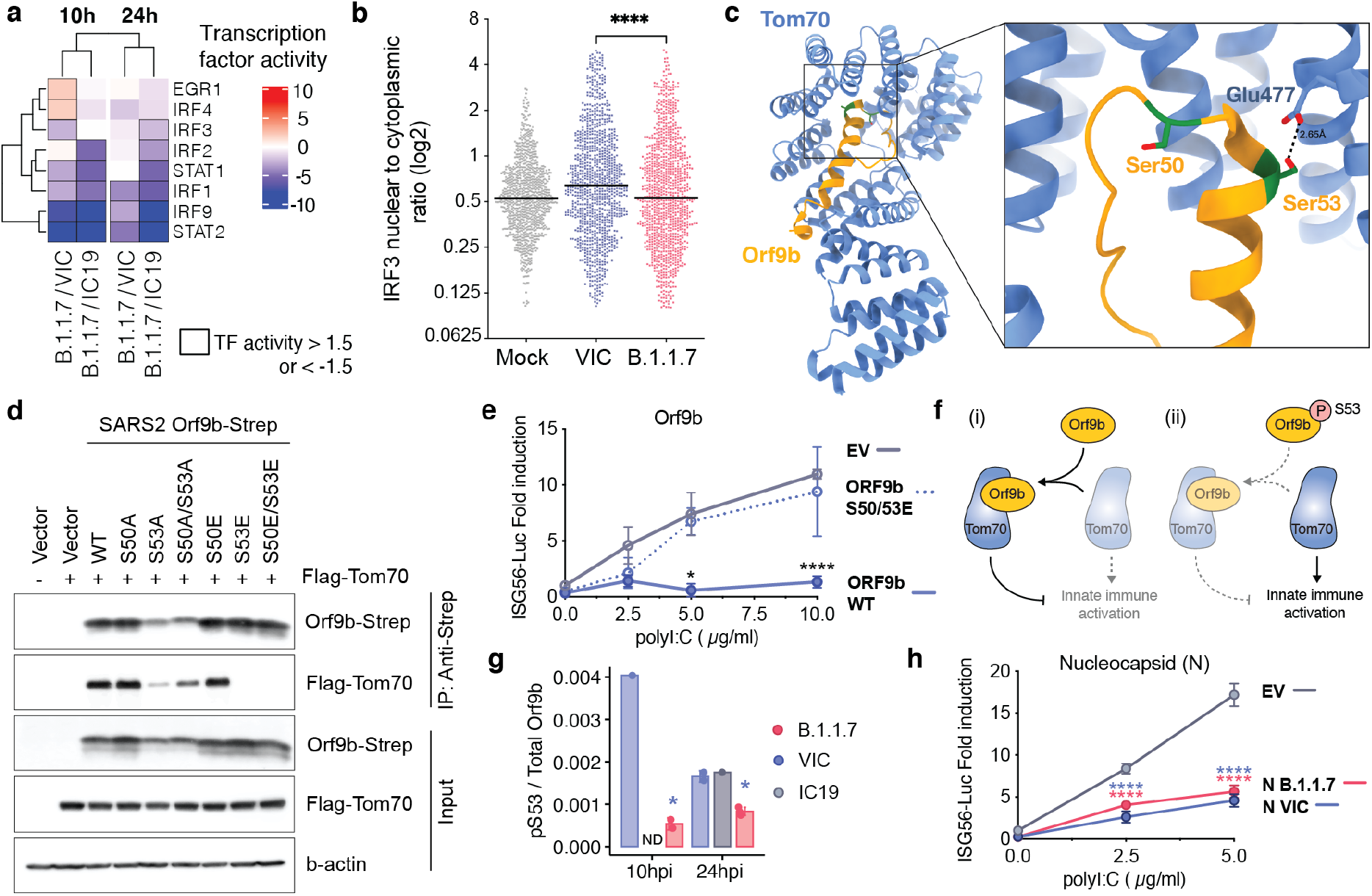
Orf9b binds TOM70 and antagonises innate immune activation downstream of RNA sensing. **a.** Transcription factor (TF) activities in the 5 top enriched terms for the RNAseq dataset (Fig. 2b, left), for each time point. TFs are clustered hierarchically along rows based on activity magnitude. Squares outlined in black depict activities > 1.5 or < −1.5. **b.** IRF3 nuclear to cytoplasmic log2 ratio in cells infected with either B.1.1.7 of VIC at an MOI of 2000 E copies/cell at 24hpi measured by single cell immunofluorescence analysis. Shown are 1000 randomly sampled cells for each condition with a cut-off of 0.1>=<5. **c.** Cryogenic electron microscopy (Cryo-EM) of SARS-CoV-2 Orf9b (yellow) in complex with TOM70 (blue) from Gordon et al. (2020b). Highlighted in red are serines (S50 and S53) in Orf9b in the TOM70 binding site. **d.** Co-immunoprecipitation of streptavidin-tagged wild-type (WT) Orf9b, and various Orf9b point mutants expressed in HEK293T cells with Flag-TOM70. Forward slash indicates the presence of both mutations. **e.** ISG56-reporter activation by poly:IC in the presence of Orf9b WT, Orf9b S50/53E or empty vector (EV) expression in HEK293T cells. **f.** Model schematic depicting proposed mechanism of innate immune antagonism by Orf9b. (i) When S53 is unphosphorylated, Orf9b binds to TOM70 and inhibits its activity in innate immune signaling. Conversely, (ii) when Orf9b is phosphorylated on S53, it can no longer interact with TOM70 and is unable to antagonise innate immune activation. **g.** Ratio between the intensity of Orf9b peptide phosphorylated on S53 and total Orf9b (as calculated in Fig. 3b, bottom) from phospho- and abundance proteomics of Calu-3 cells infected with indicated viruses for either 10 or 24 hpi (as depicted in Fig. 2a). **h.** ISG56-reporter activation by poly:IC in the presence of N (VIC), N (B.1.1.7) or empty vector (EV) expression in HEK293T cells. Statistical comparisons are performed by Mann-Whitney Test comparison (b), Two Way ANOVA with Tukey’s multiple comparison post test (e,g,h). For (e) black stars indicate the comparison between ORF9b WT and ORF9b S50/53E, For (g), blue stars indicate comparison between B.1.1.7 and VIC (blue bars). For (h), blue stars indicate comparison between VIC and EV and red stars indicate comparison between B.1.1.7 and EV. * (p<0.05), ** (p<0.01), *** (p<0.001), **** (p<0.0001).

Decreased TBK1 activity in B.1.1.7 infection (Fig. 2g) also suggests potent antagonism upstream of IRF3 by additional mechanisms. We have previously reported that SARS-CoV-2 Orf9b, which is expressed to significantly higher levels by B.1.1.7 (Fig. 3), interacts with human TOM70^41^, a mitochondrial import receptor required for MAVS activation of TBK1 and IRF3 and subsequent Type I interferon production^42^ downstream of RNA sensors. A recent study has corroborated this interaction and demonstrated inhibition of Type I interferon production by Orf9b through TOM70 interaction^39^. We previously found that two serine residues buried within the Orf9b-TOM70 binding pocket, Orf9b S50 and S53, are phosphorylated during SARS-CoV-2 infection^43–45^ (Fig. 4c). Here we discovered that mutating Orf9b S53 or S50/S53 to the phosphomimetic glutamic acid, and to a lesser extent alanine, disrupted co-immunoprecipitation of Orf9b and TOM70 (Fig 4d). Accordingly, the phosphomimetic mutations S50/53E abolished Orf9b antagonism of *ISG56-* luciferase reporter gene activation induced by poly I:C transfection that mimics RNA sensing (Fig. 4e), presumably by preventing interaction with TOM70. This suggests that Orf9b suppresses signalling downstream of MAVS by targeting TOM70 and that this process is regulated by phosphorylation (Fig. 4f). Intriguingly, we detected lower levels of B.1.1.7 Orf9b S53 phosphorylation at 10hpi compared to VIC (Fig. 4g), an effect weakened at 24hpi, in line with suppression of host kinase activity in the early stages of B.1.1.7 infection (Fig. 2f, 2g). This suggests that not only does B.1.1.7 express more Orf9b early in infection, lower kinase activation ensures maximal Orf9b innate antagonism. At later time points (24hpi), this difference in Orf9b phosphorylation is less pronounced, consistent with a modest increase in kinase activity at 24hpi for B.1.1.7 compared to first wave isolates (Fig. 2g).

SARS-CoV-2 N has also been shown to inhibit activation of RNA sensing^16^. B.1.1.7 has a modest increase in N expression (Fig. 3a, 3d), and has acquired 4 N coding changes (Fig. 1a). We therefore tested whether B.1.1.7 N displays enhanced innate immune antagonism. In fact, B.1.1.7 N antagonism of poly I:C activation of a *ISG56*-luciferase reporter was comparable to antagonism by VIC N, suggesting the coding changes do not enhance B.1.1.7 N potency (Fig. 4h).

## Discussion

Our data reveal how the SARS-CoV-2 B.1.1.7 lineage has adapted to the host by enhancing antagonism of the innate immune response. Strikingly, we find that B.1.1.7 has specifically increased subgenomic RNA synthesis and expression of key viral innate antagonists, Orf9b as well as Orf6 and nucleocapsid (N) protein (Fig. 5). This remarkable and novel observation suggests evolution of B.1.1.7 nucleotide sequences that modulate specific sgRNA production, and selection for increased sgRNA synthesis and protein expression, rather than selection of protein coding changes to alter or enhance viral antagonist function. Accordingly, we found that B.1.1.7 nucleocapsid protein coding changes did not enhance inhibition of RNA sensing. However, given SARS-CoV-2 encodes multiple, functionally overlapping innate immune antagonists^12–14^, it is possible that B.1.1.7 protein variation, for example in Nsp3 or Nsp6, could also contribute to enhanced immune antagonism. Importantly, increased detection and expression of Orf9b, Orf6 and N sgRNA has been reported in B.1.1.7 patient samples^40^, supporting the *in vivo* relevance of our findings.

**Figure 5.**
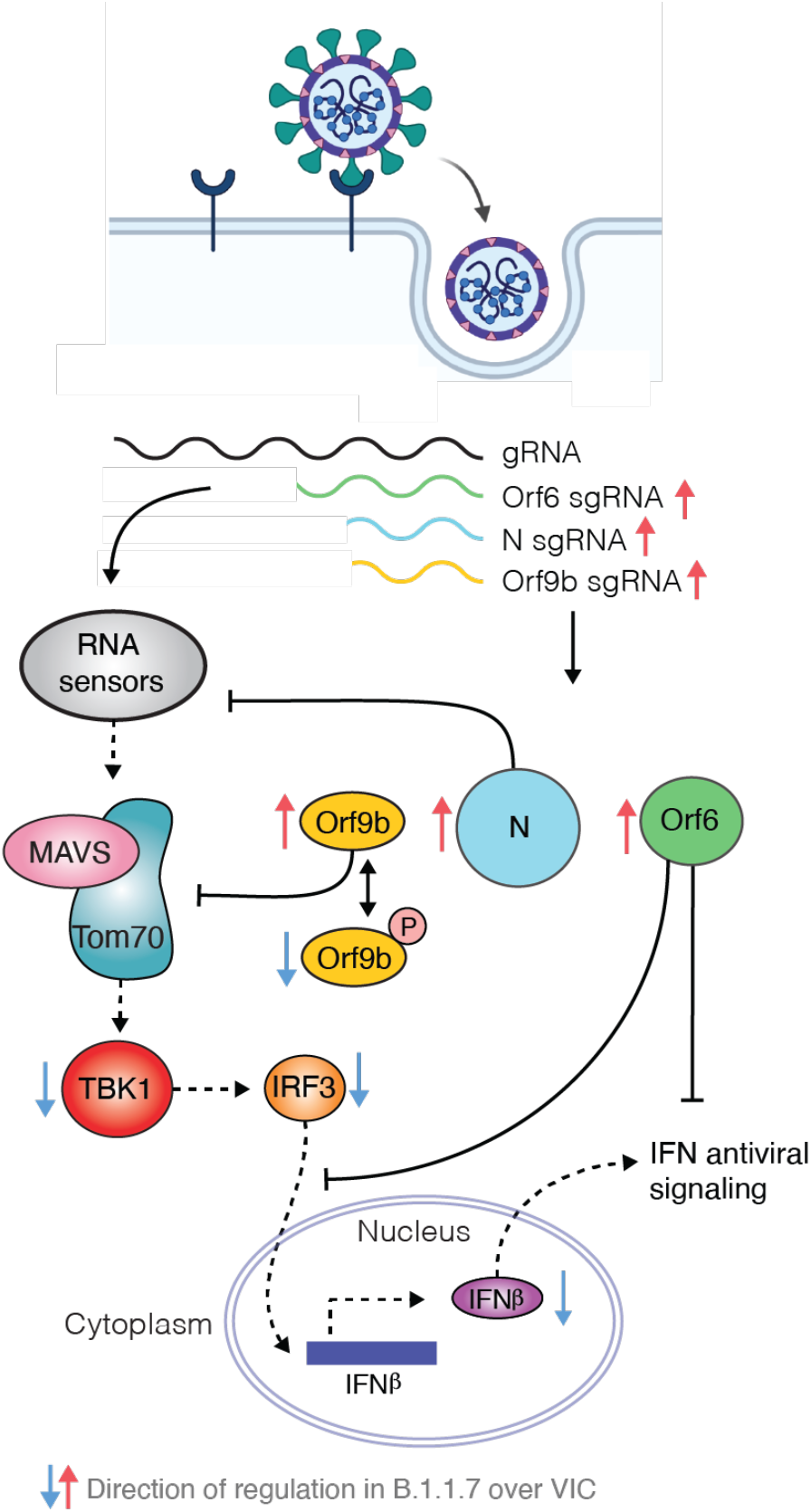
Model schematic depicting how B.1.1.7 antagonises innate immune activation. Highly transmissible SARS-CoV-2 B.1.1.7 has evolved to more effectively antagonise the innate immune response. SARS-CoV-2 wave one isolates activate a delayed innate response in airway epithelial cells relative to rapid viral replication, indicative of viral antagonism of innate immune responses early in infection. It is known that Orf9b, Orf6 and N are innate immune antagonists, acting at different levels to inhibit RNA sensing. Orf6 inhibits IRF3 and STAT1 nuclear translocation^13,24^, N prevents activation of RNA sensor RIG-I^16^ and here we show that Orf9b inhibits RNA sensing through interaction with TOM70 regulated by phosphorylation. We find that B.1.1.7 has evolved to produce more sgRNA for these key innate immune antagonists leading to increased protein levels and enhanced innate immune antagonism as compared to first wave isolates.

Coronavirus sgRNAs are produced by discontinuous transcription during negative-strand RNA synthesis, regulated by RNA elements called transcriptional regulatory sequences (TRS)^46^. Complementarity between the TRS upstream of each Orf and the TRS near the leader sequence at the 5’ end of the genome (TRS-L) mediates production of nascent sgRNAs with a 5’ leader derived from genomic RNA. Orf9b is an alternative reading frame in N, expressed from its own sgRNA. In wave one isolates, the Orf9b TRS has weak complementarity with TRS-L, consistent with low levels of Orf9b protein and sgRNA. However enhanced B.1.1.7 Orf9b sgRNA synthesis is likely mediated by changes in N (28,880 GAT>CAT, D3L) that enhance complementarity between the Orf9b TRS and TRS-L surrounding sequences^40^. At this moment, we cannot exclude whether the increase in Orf9b protein expression detected by proteomics is due to increased levels of Orf9b subgenomic RNA or to its increased translation from the N sgRNA, or to both. It is striking that Orf9b is not only enhanced in expression in B.1.1.7 but appears to be regulated by phosphorylation which is in turn particularly repressed during B.1.1.7 infection. This suggests that Orf9b inhibition of innate immunity is regulated by the host innate response itself. In this model, unphosphorylated Orf9b is maximally active early after infection to permit effective innate antagonism and viral production, but as host activation begins, Orf9b becomes phosphorylated and switched off, enabling subsequent innate immune activation. Such an inflammatory switch may have evolved by coronaviruses to enhance transmission by increasing inflammation at the site of infection once virus production is high, leading to symptoms that promote transmission such as mucosal secretions and coughing.

Detection of the N* sgRNA in B.1.1.7 infected cells can be explained by a triple nucleotide change spanning amino acids R203K/G204R (28881 GGG>AAC), which creates a novel TRS, near a downstream start codon, predicted to generate a short C-terminal form of N called N*^40,47^, which may have innate immune antagonist activity. Orf6 is known to antagonise the innate response through inhibition of transcription factor STAT1 and IRF3 nuclear entry^13,24^. Intriguingly, the increase in Orf6 sgRNA expression cannot be explained by any changes around its TRS, which has weak complementarity to the TRS-L, suggesting evolution of a different regulatory mechanism in B.1.1.7 to increase Orf6 expression. Future studies will be important to understand which genotypic adaptations confer this phenotype.

We propose that enhanced innate immune antagonism by the B.1.1.7 lineage contributes to its transmission advantage, as has been observed for HIV, another emergent pandemic virus^17,18^. We hypothesise that more effective innate immune antagonism permits enhanced transmission through reduced and delayed host responses which otherwise protect cells from infection. We propose that our model captures the earliest interactions between the virus and airway epithelial cells, in which the virus outpaces the innate response through a combination of antagonism and evasion. In the Calu-3 system, differences in innate immune antagonism between variants do not translate to differences in viral replication kinetics. We have previously shown that even for wave one isolates, the innate response occurs too late to restrict replication in Calu-3 cells^23^. We hypothesise that *in vivo*, enhanced innate antagonism could promote B.1.1.7 replication to higher levels and permit *in vivo* dissemination, in line with observations of delayed symptom onset for infections, and enhanced inflammatory disease ^5,6^. This is also consistent with reports of prolonged viral shedding of B.1.1.7^48,49^, suggesting less effective control of B.1.1.7 replication, both of which may enhance transmission.

Our data highlight that changes in protein expression levels may have significant impact on the virus-host interaction. This has important implications for management of the ongoing pandemic. It is expected that expanding ongoing sequencing efforts to monitor subgenomic RNA levels^40^ will be critical in identification of future SARS-CoV-2 variants of concern. Other reports suggest increased affinity of the B.1.1.7 spike protein for human ACE2^50^, which may enhance viral entry efficiency and therefore transmission, and is in line with B.1.1.7 adaptation to its new human host. Our findings highlight the importance of studying changes outside Spike to understand the phenotype of B.1.1.7, other current variants, and future variants, and to emphasise the importance of innate immune evasion in the ongoing process of adaptation of SARS-CoV-2 to a new host.

## Acknowledgements

This research was funded by grants from the National Institutes of Health (P50AI150476, U19AI135990, U19AI135972, R01AI143292, R01AI120694, and P01AI063302 to N.J.K.; F32CA239333 to M.B.; R01GM133981 to D.L.S.); by the Excellence in Research Award (ERA) from the Laboratory for Genomics Research (LGR), a collaboration between UCSF, UCB, and GSK (#133122P); by the Roddenberry Foundation, by funding from F. Hoffmann-La Roche and Vir Biotechnology and gifts from QCRG philanthropic donors. This research was also partly funded partly funded by CRIP (Center for Research on Influenza Pathogenesis), a NIAID funded Center of Excellence for Influenza Research and Surveillance (CEIRS, contract #HHSN272201400008C), and CRIPT (Center for Research on Influenza Pathogenesis and Transmission), a NIAID funded Center of Excellence for Influenza Research and Response (CEIRR, contract #75N93021C00014), by NCI SeroNet grant U54CA260560, and by the generous support of the JPB Foundation, the Open Philanthropy Project (research grant 2020-215611 (5384)), and anonymous donors to AG-S. This work was also supported by the Defense Advanced Research Projects Agency (DARPA) under Cooperative Agreement #HR0011-19-2-0020. The views, opinions, and/or findings contained in this material are those of the authors and should not be interpreted as representing the official views or policies of the Department of Defense or the U.S. Government.

GJT was funded by Wellcome Senior Fellowship 108183 followed by Wellcome Investigator Award 220863. CJ was funded by Wellcome Investigator Award 108079. GJT and CJ were funded by MRC/UKRI G2P-UK National Virology consortium (MR/W005611/1) and the UCL COVID-19 fund. MN was funded by Wellcome Investigator Award 207511. IG is a Wellcome Senior Fellow and this work was supported by grants from the Wellcome Trust (Refs: 207498 and 206298). Funds were also obtained from the National Institutes of Health Research UCL/UCLH Biomedical Research Centre. MVXW is supported by the NIHR Biomedical Research Centre at UCLH and IDEA Bio-Medical Ltd. We are grateful to the National Institute of Health Research Health Protection Research Unit in Respiratory Infections (NIHR #200927), the Assessment of Transmission and Contagiousness of COVID-19 in Contacts (ATACCC) Study funded by the DHSC COVID-19 Fighting Fund. We are also very grateful to the ATACCC investigators, in particular Ajit Lalvani, Jake Dunning, Joe Fenn, Rhia Kundu, Robert Varro, Sarah Hammett, Jessica Cutajaar, Eimear McDermott, Jada Samuel, Samuel Bremang, Alexandra Koycheva, Nieves Fernandez Derqui, Sam Janakan, Emily Conibear, Lulu Wang & Seran Hakki and Maria Zambon, Joanna Ellis, Angie Lackenby, Shajahan Miah and colleagues at Public Health England and Giada Mattiuzzo at the National Institute for Biological Standards and Controls and Wendy Barclay and Jonathan Brown at Imperial college London for provision of variant isolates, reagents and advice. We are grateful to Richard Milne at UCL at University of Cambridge for valuable discussions and critical reading of the manuscript.

## Author Contributions

Conceptualisation: LGT, MB, AR, LZA, CJ, GJT, NJK; Experimental setup, investigation and analysis: LGT, MB, AR, LZA, BP, MVXW, MU, AR, JT, KO, HB, MS, AR, KC, BH, DM, MH, JH, AJ, IGG, JMF, KS, NJ, KV, MN, PB, DLS, AGS, CJ, GJT, NJK; Writing, review and editing: LGT, MB, AR, LZA, BP, MVXW, MU, AR, JT, KO, HB, MS, AR, KC, BH, DM, MH, JH, AJ, IGG, JMF, KS, NJ, KV, MN, PB, DLS, AGS, CJ, GJT, NJK. Coordination and supervision: AJ, IGG, JMF, KS, NJ, KV, MN, PB, DLS, AGS, CJ, GJT, NJK.

## Materials & Correspondence

Correspondence for materials should be addressed to (NJK), (GJT), and.

## Supplemental Tables

Table S1. Fold changes and p-values for RNAseq, abundance proteomics, and phosphoproteomics datasets.

Table S2. Full pathway enrichment results of RNAseq, abundance proteomics, and phosphoproteomics datasets (i.e. Figures 2b and S1g-i).

Table S3. Fold changes and p-values for interferon stimulated genes from RNAseq and abundance proteomics datasets (i.e. Figures 2c–d).

Table S4. Full table of calculated kinase activities for comparisons between B.1.1.7, VIC, and IC19.

Table S5. Full table of calculated transcription factor activities for comparisons between B.1.1.7, VIC, and IC19.

Table S6. Viral RNA and protein quantities and ratios for B.1.1.7 to VIC and IC19 (i.e. Figure 3 and S3).

Table S7. Read counts of subgenomic RNA mapped to SARS-CoV-2 genome (i.e. Figure 3i).

## Data availability

Abundance proteomics and phosphoproteomics datasets have been deposited to the ProteomeXchange Consortium via the PRIDE partner repository with the dataset identifier PXD026302. Reviewers may access the raw data with the username and password “KBANyPDu”. Raw RNAseq data files are available from the corresponding authors upon request.

## Methods

### Cell culture

Calu-3 cells were purchased from ATCC (HTB-55) and Caco-2 cells were a kind gift from Dr. Dalan Bailey (Pirbright Institute, USA). Hela-ACE2 cells were a kind gift from Dr. James E Voss (TSRI, USA)^51^. HEK293T cells were a kind gift from Jeremy Luban. Cells were cultured in Dulbecco’s modified Eagle Medium (DMEM) supplemented with 10% heat-inactivated FBS (Labtech), 100U/ml penicillin/streptomycin, with the addition of 1% Sodium Pyruvate (Gibco) and 1% Glutamax. All cells were passaged at 80% confluence. For infections, adherent cells were trypsinised, washed once in fresh medium and passed through a 70 μm cell strainer before seeding at 0.2×10^6^ cells/ml into tissue-culture plates. Calu-3 cells were grown to 60-80% confluence prior to infection as described previously^23^.

### Viruses

SARS-CoV-2 isolate VIC was provided by NISBC, and IC19, B.1.1.7, B.1.1.7 (B) and B.1.1.7 (C) are reported in ^11^, full isolate names and GISAID references are listed below. Viruses were propagated by infecting Caco-2 cells at MOI 0.01 TCID50/cell, in culture medium at 37°C. Virus was harvested at 72 hours post infection (hpi) and clarified by centrifugation at 4000 rpm for 15 min at 4°C to remove any cellular debris. We have previously shown that infection of Caco-2 cells in these conditions does not result in activation of the innate response or cytokine carryover ^23^. Virus stocks were aliquoted and stored at −80°C. Virus stocks were quantified by extracting RNA from 100μl of supernatant with 1μg carrier RNA using Qiagen RNeasy clean up RNA protocol, before measuring viral E RNA copies per ml by RT-qCPR as described below.

| Virus ID in manuscript | Isolate name | PANGO lineage | GISAID Accession ID |
| --- | --- | --- | --- |
| VIC | BetaCoV/Australia/VIC01/2020 | B | - |
| IC19 | hCoV-19/England/IC19/2020 | B.1.13 | EPI_ISL_475572 |
| B.1.1.7 | hCoV-19/England/204690005/2020 | B.1.1.7 | EPI_ISL_693401 |
| B.1.1.7 (B) | hCoV-19/England/205090256/2020 | B.1.1.7 | EPI_ISL_747517 |
| B.1.1.7 (C) | hCoV-19/England/205080610/2020 | B.1.1.7 | EPI_ISL_723001 |

### Viral sequencing and assembly

Viral stocks were sequenced to confirm each stock was the same at consensus level to the original isolate. Sequencing was performed using a multiplex PCR-based approach using the ARTIC LoCost protocol and v3 primer set as described^52,53^. Amplicon libraries were sequenced using MinION flow cells v9.4.1 (Oxford Nanopore Technologies, Oxford, UK). Genomes were assembled using reference-based assembly to the MN908947.3 sequence and the ARTIC bioinformatic pipeline using 20x minimum coverage cut-off for any region of the genome and 50.1% cut-off for calling single nucleotide polymorphisms.

### Infection of human cells

For infections, multiplicities of infection (MOI) were calculated using E copies/cell quantified by RT-qPCR. Cells were inoculated with diluted virus stocks for 2h at 37°C, subsequently washed once with PBS and fresh culture medium was added. At indicated time points, cells were harvested for analysis.

### Virus quantification by TCID50

Virus titres were determined by 50% tissue culture infectious dose (TCID50) on Hela-ACE2 cells. In brief, 96 well plates were seeded at 5×10^3^ cells/well in 100 μl. Eight ten-fold serial dilutions of each virus stock or supernatant were prepared and 50 μl added to 4 replicate wells. Cytopathic effect (CPE) was scored at 2-3 days post infection. TCID50/ml was calculated using the Reed & Muench method, and an Excel spreadsheet created by Dr. Brett D. Lindenbach was used for calculating TCID50/mL values^54^.

### RT-qPCR of viral proteins in infected cells

RNA was extracted using RNeasy Micro Kits (Qiagen) and residual genomic DNA was removed from RNA samples by on-column DNAse I treatment (Qiagen). Both steps were performed according to the manufacturer’s instructions. cDNA was synthesised using SuperScript III with random hexamer primers (Invitrogen). RT-qPCR was performed using Fast SYBR Green Master Mix (Thermo Fisher) for host gene expression and subgenomic RNA expression or TaqMan Master mix (Thermo Fisher) for viral RNA quantification, and reactions performed on the QuantStudio 5 Real-Time PCR systems (Thermo Fisher). Viral E RNA copies were determined by a standard curve, using primers and a Taqman probe specific for E, as described elsewhere 55 and below. The primers used for quantification of viral subgenomic RNA are listed below, the same forward primer against the leader sequence was used for all reactions, and is as described by the Artic Network^40,52^. Using the 2-ΔΔCt method, sgRNA levels were normalised to GAPDH to account for differences in RNA loading and then normalised to the level of ORF1a gRNA quantified in the same way for each variant to account for differences in the level of infection. Host gene expression was determined using the 2-ΔΔCt method and normalised to GAPDH expression using primers listed below.

The following primers and probes were used:

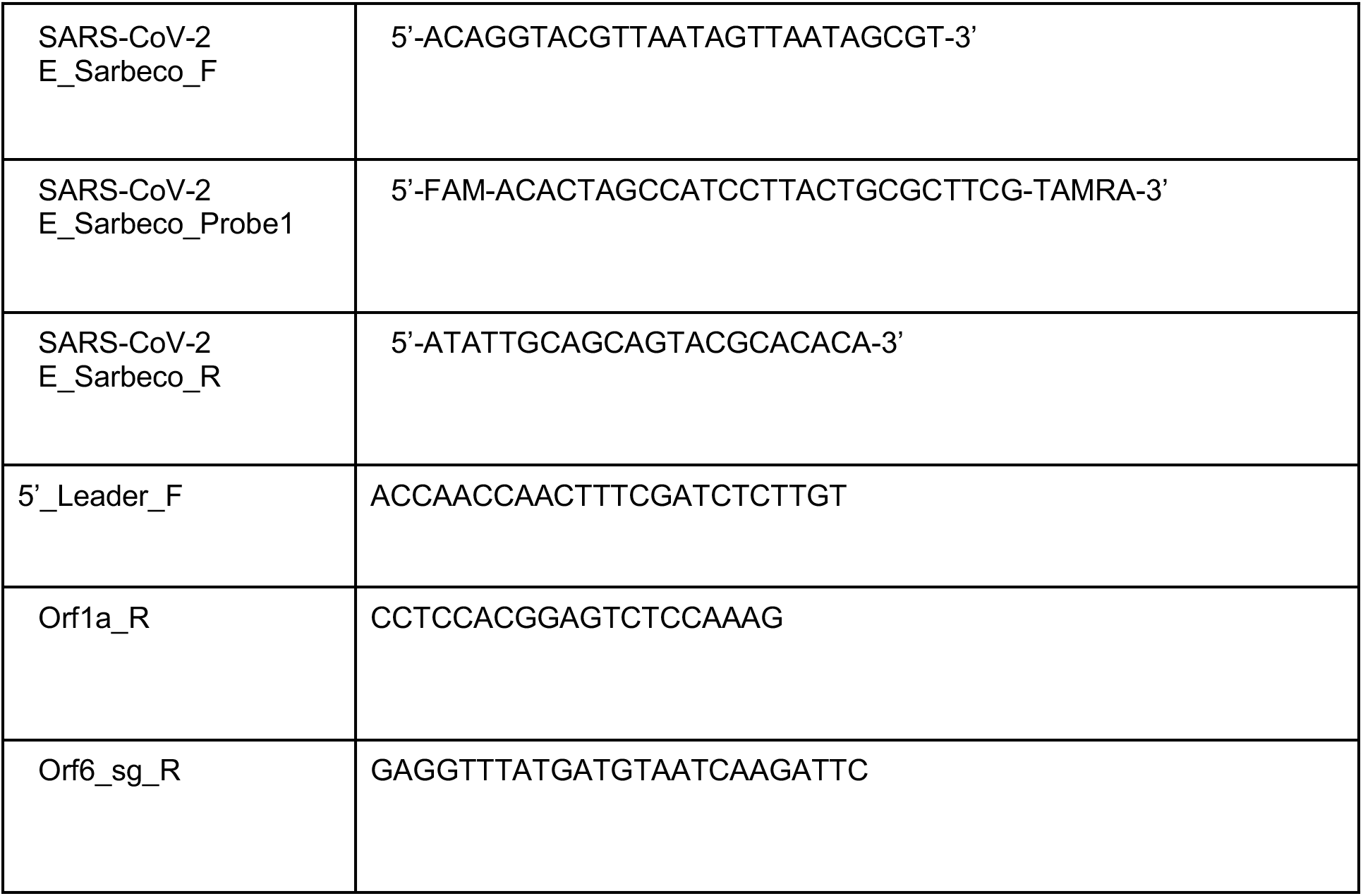
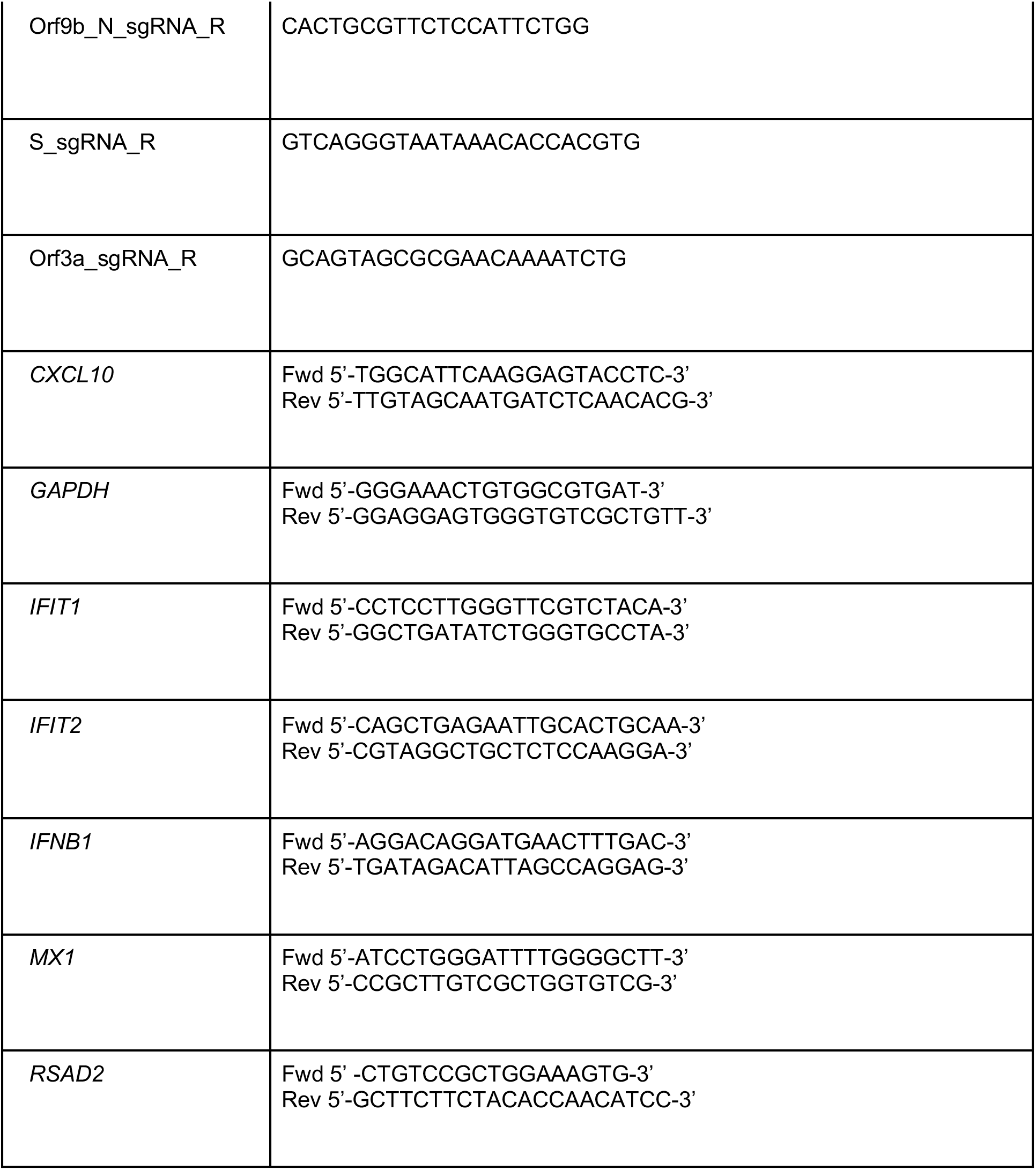

### Western blot for viral proteins in infected cells

For detection of N, Orf6, spike and tubulin expression, whole cell protein lysates were extracted with RIPA buffer, and then separated by SDS-PAGE, transferred onto nitrocellulose and blocked in PBS with 0.05% Tween 20 and 5% skimmed milk. Membranes were probed with rabbit-anti-SARS spike (Invitrogen, PA1-411-1165, 0.5ug/ml), rabbit-anti-Orf6 (Abnova, PAB31757, 4ug/ml), Cr3009 SARS-CoV-2 cross-reactive human-anti-N antibody (1ug/ml) (a kind gift from Dr. Laura McCoy, UCL), mouse-anti-alpha-tubulin (SIGMA, clone DM1A) followed by IRDye 800CW or 680RD secondary antibodies (Abcam, goat anti-rabbit, goat anti-mouse or goat anti-human). Blots were Imaged using an Odyssey Infrared Imager (LI-COR Biosciences) and analysed with Image Studio Lite software.

### Flow cytometry of infected cells

For flow cytometry analysis, adherent cells were recovered by trypsinisation and washed in PBS with 2mM EDTA (PBS/EDTA). Cells were stained with fixable Zombie UV Live/Dead dye (Biolegend) for 6 min at room temperature. Excess stain was quenched with FBS-complemented DMEM. Unbound antibody was washed off thoroughly and cells were fixed in 4% PFA prior to intracellular staining. For intracellular detection of SARS-CoV-2 nucleoprotein, cells were permeabilised for 15 min with Intracellular Staining Perm Wash Buffer (BioLegend). Cells were then incubated with 1μg/ml CR3009 SARS-CoV-2 cross-reactive antibody (a kind gift from Dr. Laura McCoy, UCL) in permeabilization buffer for 30 min at room temperature, washed once and incubated with secondary Alexa Fluor 488-Donkey-anti-Human IgG (Jackson Labs). All samples were acquired on a BD Fortessa X20 using BD FACSDiva software. Data was analysed using FlowJo v10 (Tree Star).

### Innate immune sensing assay

HEK293T cells were seeded in 48-well plates (5×10^4^ cells/well) the day before transfection. For viral protein expression, cells were transfected with 100ng of empty vector or vector encoding either ORF9b, ORF9bS50/53E, VIC N or B.1.1.7 N (pLVX-EF1alpha-IRES-Puro backbone), alongside 10ng of ISG56-firefly luciferase reporter plasmid (kindly provided by Andrew Bowie, Trinity College Dublin), and 2.5ng of a Renilla luciferase under control of thymidine kinase promoter (Promega), as a control for transfection. Transfections were performed with 0.75μL fugene (Promega) and 25μl Optimem (Gibco) per well. Cells were stimulated 24 hours post plasmid transfection with poly I:C (Invivogen), concentrations stated in figures (final 250μl volume per well), using Lipofectamine 2000 (Invitrogen) at a 3:1 ratio and 25μl optimem. Cells were lysed with 100 μl passive lysis buffer (Promega) 24 h after stimulation, 30 μl of cell lysis was transferred to a white 96-well assay plate and firefly and renilla activities were measured using the Dual-Glo® Luciferase Assay System (Promega), reading luminescence on a GloMax ®-Multi Detection System (Promega). For each condition, data were normalized by dividing the firefly luciferase activity by renilla luciferase activity and then compared to the empty-vector transfected mock-treated control to generate a fold induction.

### Immunofluorescence staining and microscopy imaging

Cells were fixed using 4% PFA-PBS for 1h and subsequently washed with PBS. A blocking step was carried out for 1h at room temperature with 10% goat serum/1%BSA in PBS. Nucleocapsid (N) protein detection was performed by primary incubation with human anti-N antibody (Cr3009, 1ug/ml) for 18h, and washed thoroughly in PBS. Where appropriate, N-protein staining was followed by incubation with mouse anti-IRF3 (sc-33641, Santa Cruz) for 1h. Primary antibodies were detected by labelling with secondary anti-human AlexaFluor-568 and anti-mouse AlexaFluor 488 conjugates (Jackson Immuno Research) for 1h. All cells were then labelled with HCS CellMask DeepRed (H32721, Thermo Fisher) and Hoechst33342 (H3570, Thermo Fisher). Images were acquired using the WiScan® Hermes High-Content Imaging System (IDEA Bio-Medical, Rehovot, Israel) at magnification 10X/0.4NA or 40X/0.75NA. Four channel automated acquisition was carried out sequentially (DAPI/TRITC, GFP/Cy5). Images were acquired at 40X magnification, 35% density/ 30% well area resulting in 102 FOV/well.

### Image analysis of immunofluorescence experiments

IRF3 raw image channels were pre-processed using a batch rolling ball background correction in FIJI imagej software package56 prior to 514 quantification. Automated image analysis was carried out using CellProfiler^57^. Firstly, Nuclei were identified as primary objects by segmentation of the Hoechst33342 channel. Cells were identified as secondary objects by nucleus-dependent segmentation of the CellMask channel. Cell cytoplasm was segmented by subtracting the nuclear objects mask from the cell masks. Nucleocapsid positive cells were identified by identifying nucleocapsid signal as primary objects followed by generation of a nucleocapsid mask which was then applied to filter the segmented cell population. Intensity properties were calculated for the nuclei, cytoplasm and cell object populations. Nuclear:cytoplasmic ratio was calculated as part of the pipeline by dividing the Integrated Intensity of the nuclei object by the integrated intensity of corresponding cytoplasm object. Plotted are 1000 randomly sampled cells selected for each condition using the ‘Pandas’ data processing package in Python 3 with a filter of 0.1>=<5.

### Coimmunoprecipitation of Tom70 with Orf9b

HEK293T were transfected with the indicated mammalian expression plasmids using Lipofectamine 2000 (Invitrogen). Twenty-four hours post-transfection, cells were harvested and lysed in NP-40 lysis buffer [0.5% Nonidet P 40 Substitute (NP-40; Fluka Analytical), 50 mM Tris-HCl, pH 7.4 at 4°C, 150 mM NaCl, 1 mM EDTA] supplemented with cOmplete mini EDTA-free protease and PhosSTOP phosphatase inhibitor cocktails (Roche). Clarified cell lysates were incubated with Streptactin Sepharose beads (IBA) for 2 hours at 4°C, followed by five washes with NP-40 lysis buffer. Protein complexes were eluted in the SDS loading buffer and were analyzed by western blotting with the indicated antibodies. Antibodies: Rabbit anti–Strep-tag II (Abcam #ab232586); Rabbit anti-beta-actin (Cell Signaling Technology #4967); Monoclonal mouse anti-FLAG M2 antibody (Sigma Aldrich, F1804), Polyclonal rabbit anti-FLAG antibody (Sigma Aldrich, F7425)

### Cell lysis and digestion for proteomics

Following the infection time course, cells in 6-well plates were washed quickly three times in ice cold 1x PBS. Next, cells were lysed in 250uL/well of 6M guanidine hydrochloride (Sigma) in 100mM Tris-HCl (pH 8.0) and scraped with a cell spatula for complete collection of the sample. Samples were then boiled for 5 minutes at 95C to inactivate proteases, phosphatases, and virus. Samples were frozen at −80C and shipped to UCSF on dry ice. Upon arrival, samples were thawed, an additional 250uL/sample of 6M guanidine hydrochloride buffer was added, and samples were sonicated for 3× for 10 seconds at 20% amplitude. Insoluble material was pelleted by spinning samples at max speed for 10 minutes. Supernatant was transferred to a new protein lo-bind tube and protein was quantified using a Bradford assay. The entire sample (approximately 600ug of total protein) was subsequently processed for reduction and alkylation using a 1:10 sample volume of tris-(2-carboxyethyl) (TCEP) (10mM final) and 2-chloroacetamide (4.4mM final) for 5 minutes at 45°C with shaking. Prior to protein digestion, the 6M guanidine hydrochloride was diluted 1:6 with 100mM Tris-HCl pH8 to enable the activity of trypsin and LysC proteolytic enzymes, which were subsequently added at a 1:75 (wt/wt) enzyme-substrate ratio and placed in a 37°C water bath for 16-20 hours. Following digestion, 10% trifluoroacetic acid (TFA) was added to each sample to a final pH ~2. Samples were desalted under vacuum using 50mg Sep Pak tC18 cartridges (Waters). Each cartridge was activated with 1 mL 80% acetonitrile (ACN)/0.1% TFA, then equilibrated with 3 × 1 mL of 0.1% TFA. Following sample loading, cartridges were washed with 4 × 1 mL of 0.1% TFA, and samples were eluted with 2 × 0.4 mL 50% ACN/0.25% formic acid (FA). 60μg of each sample was kept for protein abundance measurements, and the remainder was used for phosphopeptide enrichment. Samples were dried by vacuum centrifugation.

### Phosphopeptide enrichment for proteomics

IMAC beads (Ni-NTA from Qiagen) were prepared by washing 3× with HPLC water, incubating for 30 minutes with 50mM EDTA pH 8.0 to strip the Ni, washing 3× with HPLC water, incubating with 50mM FeCl3 dissolved in 10% TFA for 30 minutes at room temperature with shaking, washing 3× with and resuspending in 0.1% TFA in 80% acetonitrile. Peptides were enriched for phosphorylated peptides using a King Flisher Flex. For a detailed protocol, please contact the authors. Phosphorylated peptides were found to make up more than 90% of every sample, indicating high quality enrichment.

### Mass spectrometry data acquisition for proteomics

Digested samples were analysed on an Orbitrap Exploris 480 mass spectrometry system (Thermo Fisher Scientific) equipped with an Easy nLC 1200 ultra-high pressure liquid chromatography system (Thermo Fisher Scientific) interfaced via a Nanospray Flex nanoelectrospray source. For all analyses, samples were injected on a C18 reverse phase column (25 cm × 75 μm packed with ReprosilPur 1.9 μm particles). Mobile phase A consisted of 0.1% FA, and mobile phase B consisted of 0.1% FA/80% ACN. Peptides were separated by an organic gradient from 5% to 30% mobile phase B over 112 minutes followed by an increase to 58% B over 12 minutes, then held at 90% B for 16 minutes at a flow rate of 350 nL/minute. Analytical columns were equilibrated with 6 μL of mobile phase A. To build a spectral library, one sample from each set of biological replicates was acquired in a data dependent manner. Data dependent analysis (DDA) was performed by acquiring a full scan over a m/z range of 400-1000 in the Orbitrap at 60,000 resolving power (@200 m/z) with a normalised AGC target of 300%, an RF lens setting of 40%, and a maximum ion injection time of 60 ms. Dynamic exclusion was set to 60 seconds, with a 10 ppm exclusion width setting. Peptides with charge states 2-6 were selected for MS/MS interrogation using higher energy collisional dissociation (HCD), with 20 MS/MS scans per cycle. For phosphopeptide enriched samples, MS/MS scans were analysed in the Orbitrap using isolation width of 1.3 m/z, normalised HCD collision energy of 30%, normalised AGC of 200% at a resolving power of 30,000 with a 54 ms maximum ion injection time. Similar settings were used for data dependent analysis of samples used to determine protein abundance, with an MS/MS resolving power of 15,000 and a 22 ms maximum ion injection time. Data-independent analysis (DIA) was performed on all samples. An MS scan at 60,000 resolving power over a scan range of 390-1010 m/z, a normalised AGC target of 300%, an RF lens setting of 40%, and a maximum injection time of 60 ms was acquired, followed by DIA scans using 8 m/z isolation windows over 400-1000 m/z at a normalised HCD collision energy of 27%. Loop control was set to All. For phosphopeptide enriched samples, data were collected using a resolving power of 30,000 and a maximum ion injection time of 54 ms. Protein abundance samples were collected using a resolving power of 15,000 and a maximum ion injection time of 22 ms.

### Spectral library generation and raw data processing for proteomics

Raw mass spectrometry data from each DDA dataset were used to build separate libraries for DIA searches using the Pulsar search engine integrated into Spectronaut version 14.10.201222.47784 by searching against a database of Uniprot Homo sapiens sequences (downloaded February 28, 2020) and 29 SARS-CoV-2 protein sequences translated from genomic sequence downloaded from GISAID (accession EPI_ISL_406596, downloaded March 5, 2020) including mutated tryptic peptides corresponding to the variants assessed in this study. For protein abundance samples, data were searched using the default BGS settings, variable modification of methionine oxidation, static modification of carbamidomethyl cysteine, and filtering to a final 1% false discovery rate (FDR) at the peptide, peptide spectrum match (PSM), and protein level. For phosphopeptide enriched samples, BGS settings were modified to include phosphorylation of S, T, and Y as a variable modification. The generated search libraries were used to search the DIA data. For protein abundance samples, default BGS settings were used, with no data normalisation performed. For phosphopeptide enriched samples, the Significant PTM default settings were used, with no data normalisation performed, and the DIA-specific PTM site localization score in Spectronaut was applied.

### Mass spectrometry data pre-processing

Quantitative analysis was performed in the R statistical programming language (version 3.6.1, 2019-07-05). Initial quality control analyses, including inter-run clusterings, correlations, principal components analysis, peptide and protein counts and intensities were completed with the R package artMS (version 1.8.1). Based on obvious outliers in intensities, correlations, and clusterings in PCA analysis, 1 run was discarded from the protein phosphorylation dataset (IC19 24h replicate 2). Statistical analysis of phosphorylation and protein abundance changes between mock and infected runs, as well as between infected runs from different variants (e.g. Kent versus VIC) were computed using peptide ion fragment data output from Spectronaut and processed using artMS. Specifically, quantification of phosphorylation based on peptide ions were processed using artMS as a wrapper around MSstats, via functions artMS::doSiteConversion and artMS::artmsQuantification with default settings. All peptides containing the same set of phosphorylated sites were grouped and quantified together into phosphorylation site groups. For both phosphopeptide and protein abundance MSstats pipelines, MSstats performs normalisation by median equalization, imputation of missing values and median smoothing to combine intensities for multiple peptide ions or fragments into a single intensity for their protein or phosphorylation site group, and statistical tests of differences in intensity between infected and control time points. When not explicitly indicated, we used defaults for MSstats for adjusted p-values, even in cases of N = 2. By default, MSstats uses Student’s t-test for p-value calculation and Benjamini-Hochberg method of FDR estimation to adjust p-values.

### Refining and filtering phosphorylation and abundance data

MSstats phosphorylation results had to be further simplified to effects at single sites. The results of artMS/MSstats are fold changes of specific phosphorylation site groups detected within peptides, so one phosphorylation site can have multiple measurements if it occurs in different phosphorylation site groups. This complex dataset was reduced to a single fold change per site by choosing the fold change with the lowest p-value, favoring those detected in both conditions being compared (i.e. non-infinite log2 fold change values). This single-site dataset was used as the input for kinase activity analysis and enrichment analysis. Protein abundance data was similarly simplified when a single peptide was mapped to multiple proteins; that is, by choosing the fold change with the lowest p-value, favoring those detected in both conditions being compared (see Table S1 for final refined data).

### RNA quality control

Thirty total RNA samples were submitted for RNA quality control. Total RNA samples were run on the Agilent Bioanalyzer, using the Agilent RNA 6000 Nano Kit. Three samples were excluded from library preparation due to severe degradation and/or low amounts of RNA present.

### Library preparation for RNAseq

Twenty-seven total RNA samples were processed using the Illumina Stranded Total RNA w/Ribo-Zero Plus assay. One-hundred nanograms of each total RNA sample (quantitated on the Invitrogen Qubit 2.0 Fluorometer using the Qubit RNA HS Assay Kit) was subjected to ribosomal RNA (rRNA) depletion through an enzymatic process, which includes reduction of human mitochondrial and cytoplasmic rRNAs. Following rRNA depletion and purification, RNA was primed with random hexamers for first-strand cDNA synthesis, then second-strand cDNA synthesis. During second-strand cDNA synthesis, deoxyuridine triphosphate (dUTP) was incorporated in place of deoxythymidine triphosphate (dTTP) to achieve strand specificity in a subsequent amplification step. Next, adenine (A) nucleotide was added to the 3’ ends of the blunt fragments to prevent ends from ligating to each other. The A-tail also provides a complementary overhang to the thymine (T) nucleotide on the 3’ end of the adapter. During adapter ligation and amplification, indexes and adapters were added to both ends of the fragments, resulting in 10bp, dual-indexed libraries, ready for cluster generation and sequencing. The second-strand was quenched during amplification due to the incorporation of dUTP during second-strand cDNA synthesis, allowing for only the antisense strand to be sequenced in Read 1. Thirteen cycles of amplification were performed.

### Library quality control and quantification for RNAseq

Each library was run on the Agilent Bioanalyzer, using the Agilent High Sensitivity DNA Kit, to assess the size distribution of the libraries. They were quantitated by quantitative polymerase chain reaction (qPCR) using a Roche KAPA Library Quantification Complete Kit (ABI Prism), and run on the Applied Biosystems QuantStudio 5 Real-Time PCR System.

### Sequencing for RNAseq

Each library was normalised to 10nM, then pooled equimolarly for a final concentration of 10nM. Pooled libraries were submitted to the University of California San Francisco Center for Advanced Technology (UCSF CAT) for one lane of sequencing on the Illumina NovaSeq 6000 S4 flow cell. The run parameter was 100×10×10×100bp.

### Viral RNA quantification from RNASeq Dataset

Viral RNA were characterised by the junction of the leader with the downstream subgenomic sequence. Reads containing possible junctions were extracted by filtering for exact matches to the 3’ end of the leader sequence “CTTTCGATCTCTTGTAGATCTGTTCTC” using the bbduk program in the BBTools package (BBTools -Bushnell B. - sourceforge.net/projects/bbmap/). This subset of leader-containing reads were left-trimmed to remove the leader, also using bbduk. The filtered and trimmed reads were matched against SARS2 genomic sequence with the bbmap program from BBtools with settings (maxindel=100, strictmaxindel=t, local=t). The left-most mapped position in the reference was used as the junction site. All strains were mapped against a reference SARS-Cov-2 sequence (accession NC_045512.2), except B.1.1.7 was mapped against a B.1.1.7-specific sequence (GISAID: EPI_ISL_693401) and the resultant positions adjusted to the reference based on a global alignment. Junction sites were labeled based on locations of TRS sequences, or other known site with a +/- 5 base pair window as follows (genomic = 67, S = 21553, orf3 = 25382, E = 26237, M = 26470, orf6 = 27041, orf7 = 27385, orf8 = 27885, N = 28257, orf9b = 28280, N* = 28878). Junction reads were counted per position, a pseudocount of 0.5 was added at all positions, counts between replicates and strains were normalised to have equal “genomic” reads, and counts were averaged across replicate samples. Means and standard errors of counts averaged across replicates were subsequently calculated. To calculate the ratios between B.1.1.7 and VIC, counts averaged across replicates from B.1.1.7 were divided in a condition and time point matched fashion by values from VIC or IC19. The standard error (se) of the ratios was calculated as (A/B) * sqrt((se.A/A)^2^ + (se.B/B)^2^).

### Host RNA analysis

All reads were mapped to the human host genome (ensembl 101) using HISAT2 aligner^58^. Host transcript abundances were estimated using human annotations (ensembl 101) using StringTie^59^. Differential gene expression were done on read counts extracted for each protein coding gene using featureCount and significance was determined by the DESeq2 R package^60^.

### Viral protein quantification

Median normalized peptide feature (peptides with unique charge states and elution times) intensities (on a linear scale) were refined to the subset that mapped to SARS-CoV-2 protein sequences using Spectronaut (see Methods). Peptide features found in the same biological replicate (i.e. due to different elution times, for example) were averaged. Next, for each timepoint separately, we selected the subset of peptides that were consistently detected in all biological replicates across all conditions (no missing values), isolating the set of peptides with the best comparative potential. We then summed all peptides mapping to each viral protein for each time point separately which resulted in our final protein intensity per viral protein per time point per biological replicate. Resulting protein intensities were averaged across biological replicates and standard errors were calculated for each condition. To calculate the ratios between B.1.1.7 and VIC, averaged intensities for B.1.1.7 were divided in a condition and time point matched fashion by values from VIC or IC19. The standard error (se) of the ratios was calculated as (A/B) * sqrt((se.A/A)^2^ + (se.B/B)^2^).

### Kinase activity analysis of phosphoproteomics data

Kinase activities were estimated using known kinase-substrate relationships in literature^61^. The resource comprises of a comprehensive collection of phosphosite annotations of direct substrates of kinases obtained from six databases, PhosphoSitePlus, SIGNOR, HPRD, NCI-PID, Reactome, and the BEL Large Corpus, and using three text-mining tools, REACH, Sparser, and RLIMS-P. Kinase activities were inferred as a Z-score calculated using the mean log2FC of phosphorylated substrates for each kinase in terms of standard error (Z = [M - u] / SE), comparing fold changes in phosphosite measurements of the known substrates against the overall distribution of fold changes across the sample. A p-value was also calculated using this approach using a two-tailed Z-test method. This statistical approach has been previously shown to perform well at estimating kinase activities^38,62^. We collected substrate annotations for 400 kinases with available data. Kinase activities for kinases with 3 or more measured substrates were considered, leaving us with 191 kinases with activity estimates in at least one or more infection time points. Kinases were clustered based on pathway similarity by constructing a kinase tree based on co-membership in pathway terms (from CP “Canonical Pathways” MSigDBv7.1).

### Pathway enrichment analysis

The pathway gene sets were obtained from the CP (i.e. “Canonical Pathways”) category of Molecular Signature Database (MSigDBv7.1)^35^. We used the same approach for this pathway enrichment analysis as we used for the kinase activity analysis. Namely, we inferred pathway regulation as Z-score and an FDR-corrected (0.05) p-value calculated from a Z-test (two-tailed) comparing fold changes in phosphosite, protein abundance, or RNA abundance measurements of genes designated for a particular pathway against the overall distribution of fold changes in the sample. All resulting terms were further refined to select non-redundant terms by first constructing a pathway term tree based on distances (1-Jaccard Similarity Coefficients of shared genes in MSigDB) between the terms. The pathway term tree was cut at a specific level (h = 0.8) to identify clusters of non-redundant gene sets. For results with multiple significant terms belonging to the same cluster, we selected the most significant term (i.e. lowest adjusted p-value). Next, we filtered out terms that were not signifƒicant (FDR corrected p-value < 0.05) for at least one contrast. Terms were ranked according to either the absolute value z-score across contrasts that included B.1.1.7 (see Fig. S1g–i) or by averaged −log10(p-values) across time-matched contrasts involving B.1.1.7 (see Fig. S2b).

### Transcription factor activity analysis

Transcription factor activities were estimated from RNAseq data using DoRothEA^63^ which provides a comprehensive resource of TF-target gene interactions and annotations indicating confidence level for each interaction based on the number of supporting evidence. We restricted our analysis to A, B, and C levels which comprise of the most reliable interactions. For the TF activity enrichment analysis, VIPER^64^ was executed with the t-statistic derived from the differential gene expression analysis between variant infected and controls (wild-type) infected cells. Transcription factor activity is defined as the normalised enrichment scores (NES) derived from the VIPER algorithm. VIPER algorithm was run with default parameters except for the eset.filter parameter which was set to FALSE and consider regulons with at least five targets.

### Selection of interferon stimulated genes (ISGs)

Interferon stimulated genes (ISGs) were taken from a prior experimental study^36^ and annotated as ISGs. To this list of 38 genes, we added the following based on manual curation from the literature: IFI16, IFI35, IFIT5, LGALS9, OASL, CCL2, CCL7, IL6, IFNB1, CXCL10, and ADAR.

## Supplemental Figures

**Figure S1.**
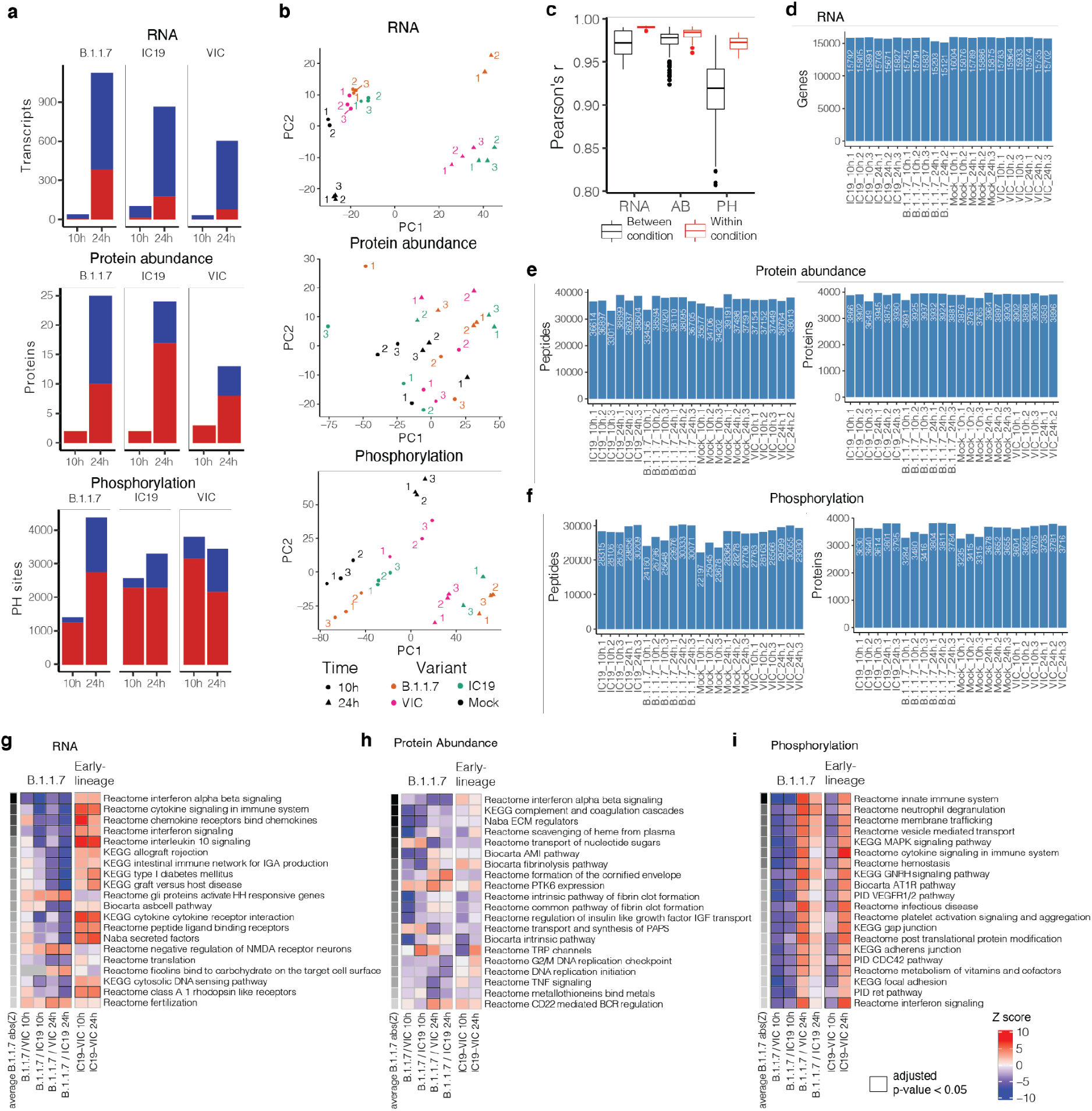
Omics data quality control and pathway enrichments. **a.** Significantly changing genes for RNA, proteins for protein abundance, and phosphorylation sites for phosphoproteomics data. Significance was defined as abs(log2FC)>1 and adjusted p-value<0.05. Red depicts positive log2 fold changes whereas blue depicts negative log2 fold changes. **b.** Principal components analysis (PCA) on normalised RNA transcripts per million (TPM), protein intensities, or phosphorylation site intensities. Non-finite values were removed and detections (transcripts, proteins, or phosphorylation sites) not shared (non-finite) between all conditions were discarded prior to analysis. **c.** Pairwise Pearson’s correlation between RNA, protein, or phosphorylation site abundance among replicates within the same condition (red) or between distinct conditions (black). **d.** Number of genes expressed above baseline in RNAseq dataset per replicate. **e.** Number of peptides and proteins detected per replicate in the abundance proteomics dataset. **f.** Number of phosphorylated peptides and corresponding proteins from phosphoproteomics dataset. **g.** Gene set enrichment analysis based on log2FC method using RNA dataset (as in Fig. 2b, see Methods). Ranking is based on the average of the absolute value z-scores across the indicated contrasts involving B.1.1.7 (per row). Enrichments with an adjusted p-value<0.05 are indicated with a black border. **h.** Same as in g, but for abundance proteomics dataset. **i.** Same as in g, but for phosphoproteomics dataset. If a protein possessed multiple phosphorylation sites, the maximum absolute value log2FC was used as the representative value for the protein. Finite values (non-infinite) were prioritised over quantitative values.

**Figure S2.**
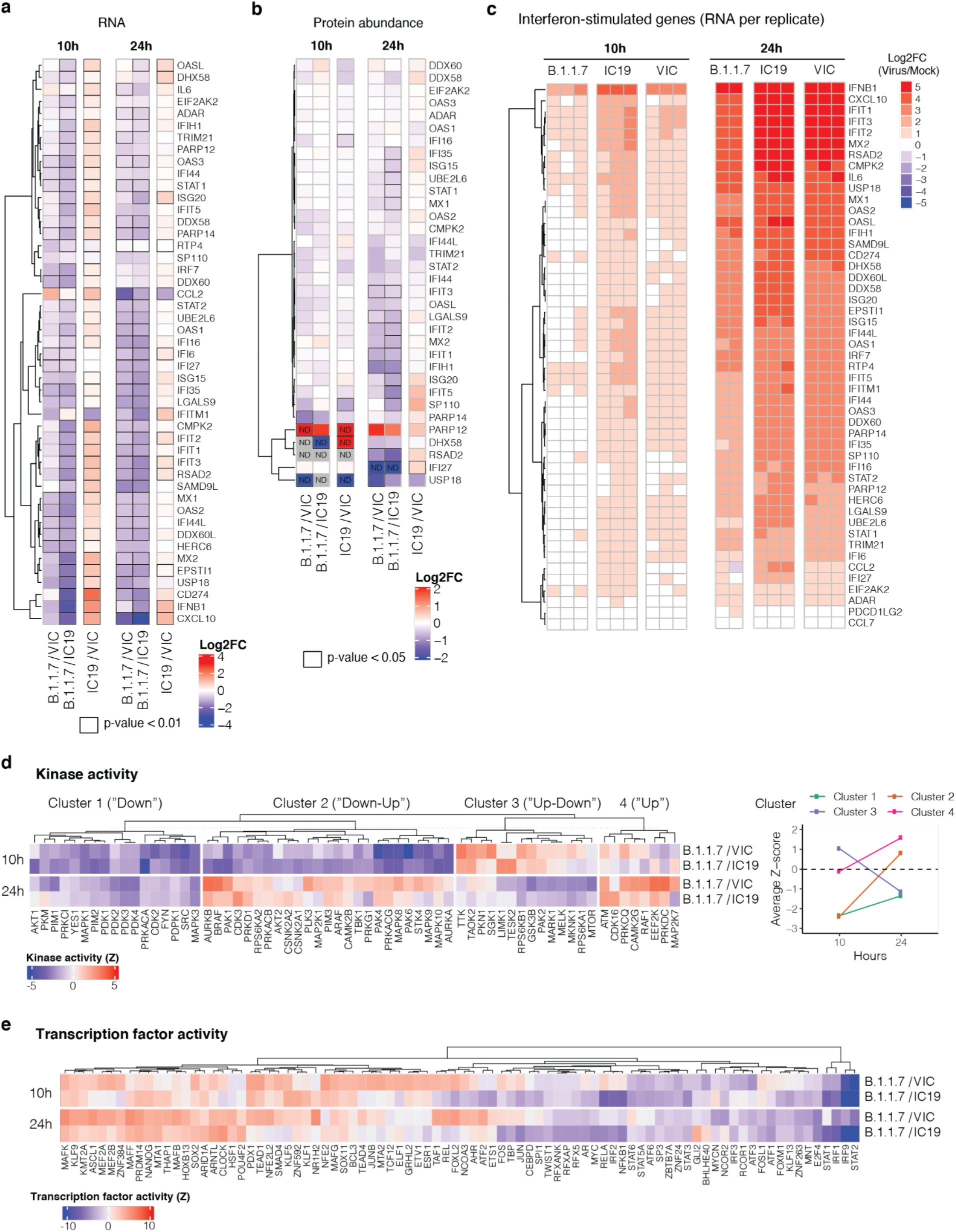
Omics data highlights recruitment of innate immune signaling. **a.** Expression of interferon-stimulated genes from Lui et al (2018)^36^ (see Methods) using the RNAseq dataset. Significant fold changes with an adjusted p-value<0.05 are indicated with black borders. **b.** Same as in (a) using the abundance proteomics dataset. N.D. indicates proteins either not detected in one condition (thus, Inf or -Inf) or not detected in both conditions. **c.** RNA expression per biological replicate of interferon-stimulated genes (ISGs) for each virus versus mock. **d.** Full kinase activity analysis of indicated contrasts. Only kinases with an absolute value z-score>2 were kept. Kinases were separated into four distinct clusters using k-means clustering, which naturally reveals groups depicting kinases downregulated for the entire time course (“Down”), downregulated early and upregulated late (“Down-Up”), upregulated early and downregulated late (“Up-Down”), or upregulated or constant throughout the time course (“Up”). Panel on right depicts the average Z-score for each distinct cluster per time point, collapsing across B.1.1.7/VIC and B.1.1.7/IC19 comparisons. **e.** Transcription factor (TF) activities were estimated from the RNAseq dataset using known TF-target gene interactions (see Methods). Only transcription factors with an absolute value NES>2.5 were kept. TF are clustered using ward hierarchical clustering based on similar activity patterns across time.

**Figure S3.**
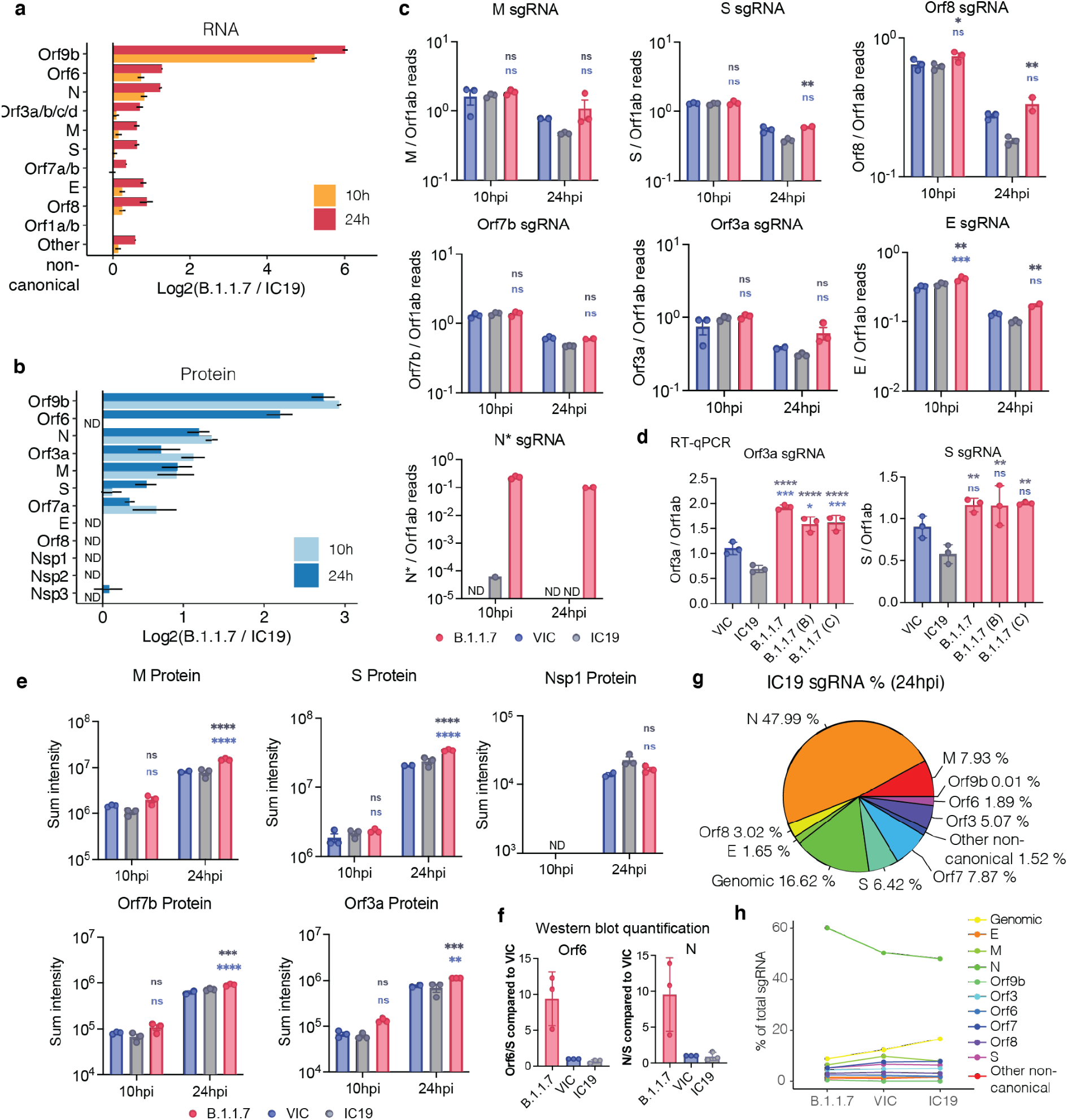
Expression of viral RNA and protein for SARS-CoV-2 variants. **a.** Log2 ratio of B.1.1.7 to IC19 subgenomic RNA (sgRNA) abundance as determined from the RNAseq dataset. sgRNA reads are counted only if they possess a leader sequence and normalised to total genomic RNA per time point and virus (see Methods). **b.** Log2 ratio of B.1.1.7 to IC19 viral proteins quantified as determined from the abundance proteomics dataset. Peptide intensities are summed per viral protein. Only peptides detected in both B.1.1.7 and IC19 are used for quantification. Bars depict the mean of three biological replicates. Error bars depict the standard error. **c.** Quantification of sgRNAs for M, S, Orf8, Orf7a, Orf3a, E and N* from the RNAseq dataset. Counts are normalised to genomic RNA abundance at each time point and virus. **d.** Quantification of Orf3a (left) or S (right) sgRNA abundance via RT-qPCR in distinct B.1.1.7 isolates, VIC, or IC19. **e.** Summed peptides per viral protein for M, S, Nsp1, Orf7b, and Orf3b from the abundance proteomics dataset. **f.** Quantification of Orf6 and N protein from western blot in Figure 3f for B.1.1.7, VIC, and IC19. **g.** Pie chart depicting proportion of total sgRNA mapping to each viral sgRNA (containing leader sequence) for IC19. **h.** Comparison of percentages of total sgRNA mapping to each viral sgRNA across B.1.1.7, VIC, and IC19. * (p<0.05), ** (p<0.01), *** (p<0.001), **** (p<0.0001). ns: non-significant.

**Figure S4.**
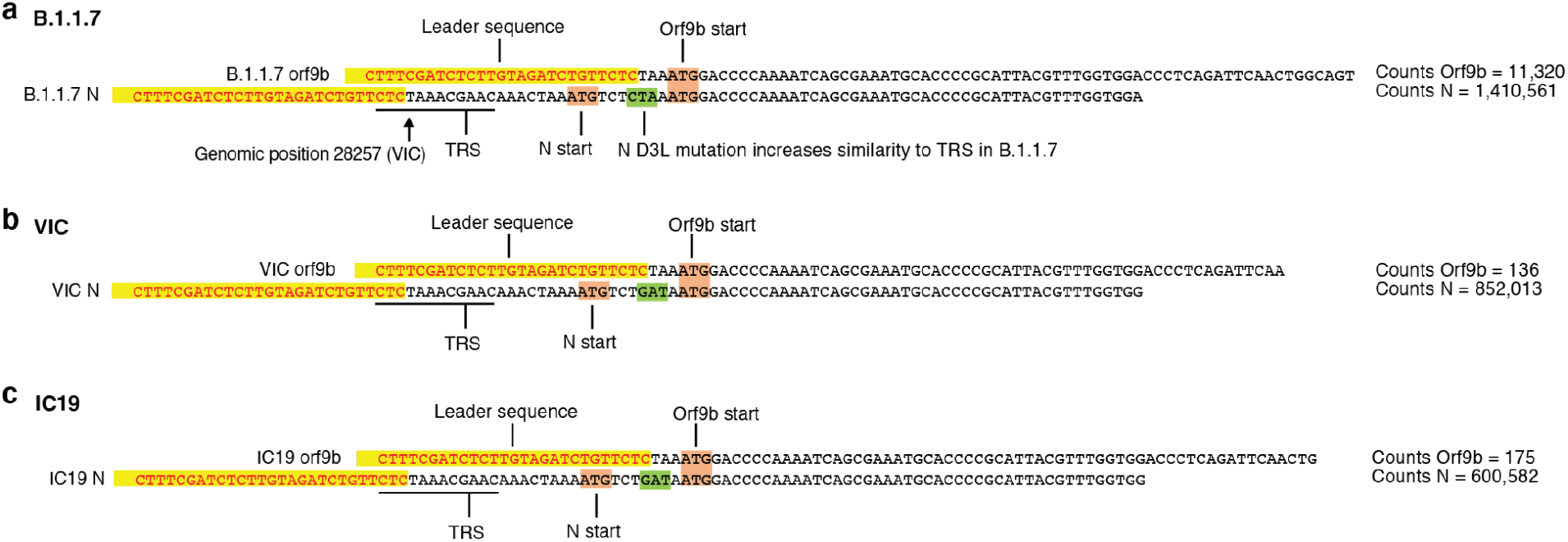
Examples of leader-containing reads for Orf9b and N from RNAseq dataset. **a.** Representative sequence for Orf9b (top) and N (bottom) sgRNA from B.1.1.7. Leader sequences used in this analysis to identify sgRNAs are highlighted in yellow. The sequence following the leader sequence is used to differentiate Orf9b versus N sgRNAs. Orf9b and N start codons are indicated in maroon. The site of the N-protein D3L mutation is indicated in green, which results in increased similarity to the transcriptional regulatory sequence (TRS) for B.1.1.7. Read counts of Orf9b and N are indicated to the right. Counts are normalized to mean genomic reads per replicate. **b.** Same as in a but for VIC. **c.** Same as in a but for IC19.

